# Precision super-resolution cryo-correlative light and electron microscopy for rapid *in situ* structural analyses of optogenetically-positioned organelles

**DOI:** 10.1101/2022.11.22.516823

**Authors:** G.M.I. Redpath, J. Rae, Y. Yao, J. Ruan, M.L. Cagigas, R. Whan, E.C. Hardeman, P.W. Gunning, V. Ananthanarayanan, R.G. Parton, N.A. Ariotti

**Affiliations:** EMBL Australia Node in Single Molecule Science, School of Medical Sciences, University of New South Wales Sydney, New South Wales, Australia; The University of Queensland, Institute for Molecular Bioscience, Brisbane, Queensland, Australia; University of New South Wales Sydney, Electron Microscope Unit, Mark Wainwright Analytical Centre, Sydney, New South Wales, Australia; University of New South Wales Sydney, School of Medical Sciences, Kensington, Sydney, New South Wales, Australia; University of New South Wales Sydney, Katharina Gaus Light Microscopy Facility, Mark Wainwright Analytical Centre, Sydney, New South Wales, Australia; The University of Queensland, Centre for Microscopy and Microanalysis, Brisbane, Queensland, Australia

## Abstract

Unambiguous targeting of cellular structures for *in situ* cryo-electron microscopy in the heterogeneous, dense, and compacted environment of the cytoplasm remains challenging. Here we have developed a novel cryogenic correlative light and electron microscopy (cryo-CLEM) workflow which combines thin cells grown on a mechanically defined substratum to rapidly analyse organelles and macromolecular complexes in the cell by cryo-electron tomography (cryo-ET). We coupled these advancements with optogenetics to redistribute perinuclear-localised organelles to the cell periphery for cryo-ET. This reliable and robust workflow allows for fast *in situ* analyses without the requirement for cryo-focused ion beam milling. We have developed a protocol where cells can be frozen, imaged by cryo-fluorescence microscopy and ready for batch cryo-ET within a day.

## Main

Advances in structural biology have seen a dramatic switch over the last 15 years from X-ray crystallography-based methods to single particle cryo-EM-based techniques and now, increasingly to structural analyses in the cellular context. *In situ* structural biology can not only provide the near atomic resolution of proteins/complexes of interest but has the added advantage of providing the context to how these proteins and macromolecular complexes interact with cellular machinery and organelles of interest; this can better inform cellular function^1-3^. While these latest advancements show almost unending possibilities for understanding how proteins and complexes elicit their cellular roles, especially in the context of visual proteomics^3^, the application of these methods have been somewhat slowed by several complicating and contributing factors. These factors include i) the intricate and expensive interconnected workflows and manual handling steps required between disparate instruments that often results in damage to the sample and ice contamination; ii) the highly concentrated environment of the cytoplasm that obscures the regions/structures/proteins of interest; and iii) the low signal to noise ratio of cryo-electron microscopy and cryo-electron tomographic datasets. As a consequence of this, the native state cellular architecture remains largely unexplored. Improvements to the ease of data collection, sample scalability and region of interest targeting will expedite the use of these techniques to the broader scientific community^1-4^.

Cryo-CLEM combines the resolving power of cryo-EM with the specific illumination of structures of interest by cryo-fluorescence microscopy (cryo-fLM) for the precise localisation of distinct cellular targets. Recent advancements in cryogenic light microscopy stages are now facilitating the combinatorial power of these previously disparate techniques. While there has been an increase in the adoption of cryo-CLEM-based methods in recent years^1,4-8^, these techniques remain an emerging field of application. Improved and integrated systems as well as simplified workflows will further facilitate the adoption of these technologies.

Here we develop a novel system for simplified correlation of structures of interest by coupling super-resolution Airyscan cryo-confocal microscopy and optogenetic manipulation techniques. We characterise PtK_2_ cell morphology and cell substate interactions to optimise and improve the yield of cryo-ET compatible regions and develop expedited cryo-fLM data acquisition strategies that deliver a simplified correlation workflow for high-precision targeting of regions of interest for *in situ* structural analyses. Finally, we develop and optimise an optogenetic-based system for analysis of perinuclear localised structures in PtK_2_ cells without the requirement for cryo-FIB. We redistribute Lysosomal Associated Membrane Protein 1 (LAMP1) from thick perinuclear regions to the thin cell boundary using blue-light stimulation and Cryptochrome2/CIBN conditional association with the kinesin motor KIF-5a for nanometre-resolution *in situ* cryo-electron tomography of late endosomes/lysosomes (here termed endolysosomes). These advancements improve the precision and broaden the applicability of cryo-CLEM for imaging of specific structures of interest in the cell. This method eliminates the requirement for Cryo-FIB as an intermediate step to mill cryo-lamellae on grids and thus i) reduces and simplifies existing manual handling steps and cryo-transfer requirements, ii) improves cryo-ET throughput and scalability and, iii) enhances the cost-effectiveness and accessibility of *in situ* cryo-EM workflows. This novel technique, which can proceed from the living cell to cryo-electron tomography of cellular organelles in one day, brings *in situ* cryo-EM within reach of all cell biologists.

## Results

We sought to develop an efficient cellular cryo-CLEM workflow that would allow for precision targeting of regions of interest by cryo-ET for cells grown on a mechanically defined substratum where organelles/structures of interest could be analyzed at high resolution without cryo-lamella preparation (Fig 1A and B). High-quality cryo-tomography requires ∼300 (or below) nm thick sample. To therefore generate cryo-tomograms without cryo-FIB milling we require a cell type with a protracted and thin cytoplasm. Towards this aim we performed a screen of multiple cell lines to determine the most appropriate cell type for *in situ* cryo-EM. We analysed general cellular morphology and cytoplasmic thickness by ultrathin vertical sections of conventionally processed cells by transmission electron microscopy (TEM). We observed that PtK_2_ cells, and to a lesser extent U-2OS and WI-38 cells, possessed a thin cytoplasm with abundant organelles extending from perinuclear regions to the cell perimeter compared to other screened lines (MDCK, Caco-2, BHK, HeLa, MEF and WI-38s; Fig 1A and B). As PtK_2_ cells were also readily transfected we chose to pursue their further characterisation.

**Figure 1.**
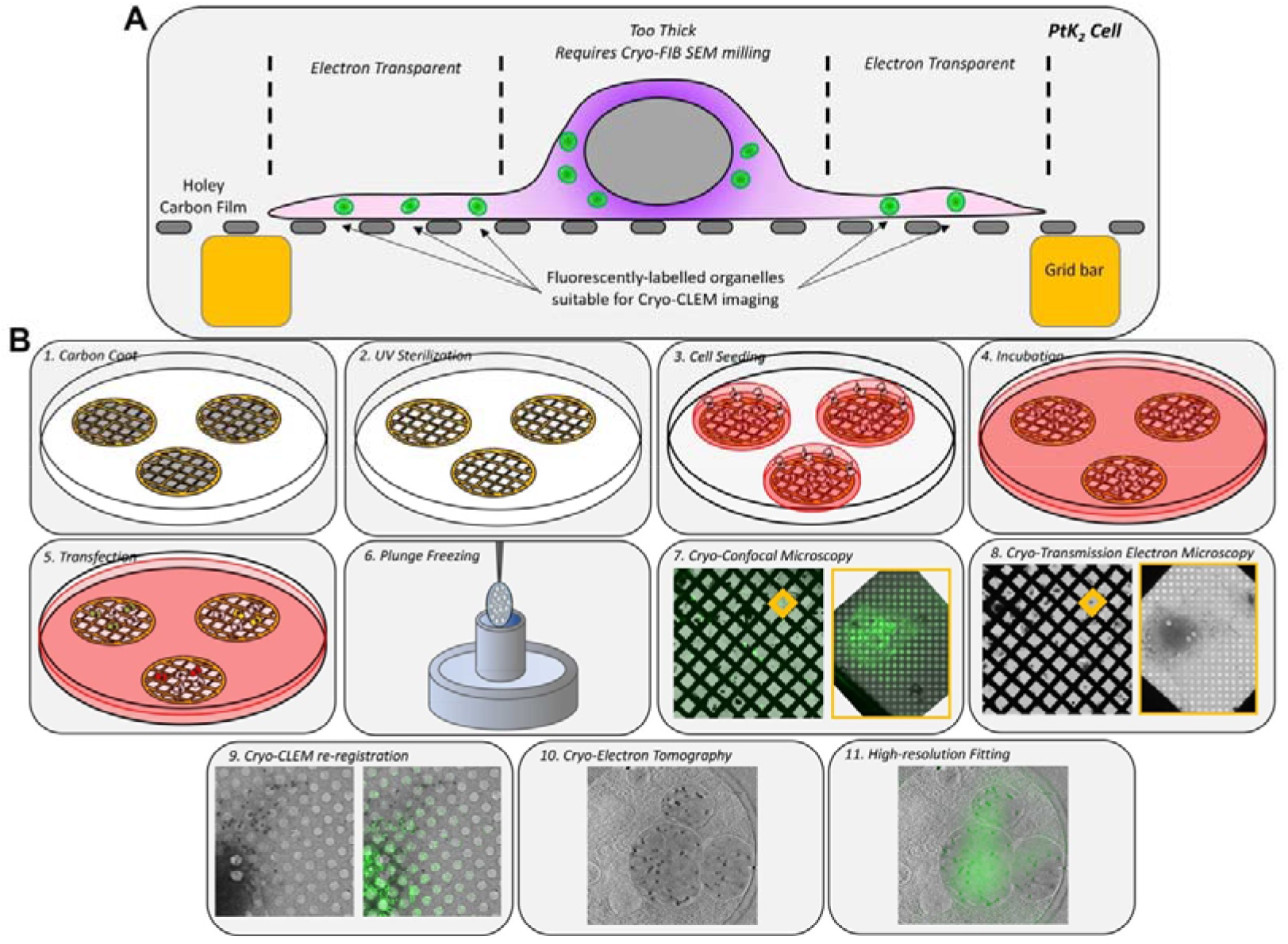
Precision super-resolution cryo-CLEM. A) Schematic of a PtK_2_ cell grown on a Quantifoil grid for the rapid *in situ* cryo-CLEM/cryo-ET. Dark purple = no electron transparency, light purple = electron transparency, green = fluorescently labelled structures of interest, grey = nucleus. B) Workflow of precision Cryo-CLEM targeting of structures of interest for *in situ* cryo-EM.

### PtK_2_ Cell-Grid substrate optimisation

*In situ* analyses require that cells can grow on grids for cryo-EM imaging and still maintain a thin cellular periphery. We therefore characterised PtK_2_ cells grown on R2/2 gold support carbon film Quantifoil grids. We performed serial blockface scanning electron microscopy (SBF-SEM) and conventional transmission electron microscopy on cells grown on grids. SBF-SEM demonstrated that PtK_2_ cells adopt a ‘fried-egg’ morphology on grids with thick nuclear and perinuclear regions that reduce into a thin cytoplasm of approximately 100-300 nm extending to the cell perimeter (Movie S1). This region contains abundant cellular organelles (Fig 2B and C; dotted circles) and cytoskeletal elements (Fig 2C’-C’’). Segmentation of cells grown on the Quantifoil grids demonstrates numerous regions of a suitable thickness for cryo-ET without the requirement for cryo-FIB lamella preparation (Fig 2C and D).

**Figure 2.**
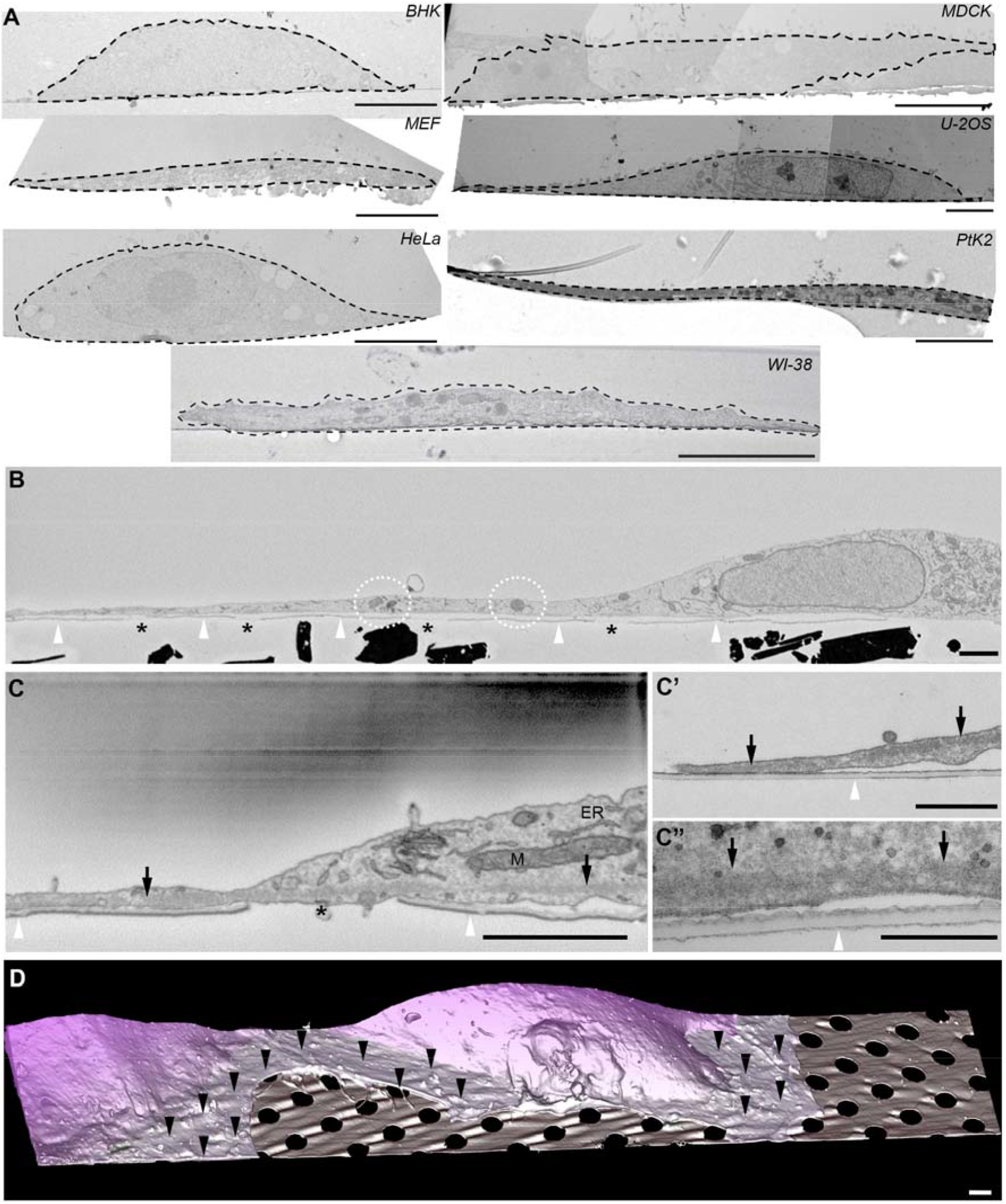
EM characterisation and morphology of PtK_2_ cells grown on Quantifoil grids. A) TEM-based ultrastructural screen of cell type thickness. 7 cell types were screened for vertical thickness. Scale = 5 µm. B) Serial block face scanning electron micrograph of a vertically oriented and resin embedded PtK_2_ cell showing a thin and extended cell periphery containing organelles (white dotted regions). White arrowheads = Quantifoil carbon film; black asterisks = holes in the Quantifoil carbon film. Scale = 2 µm. C) Intermediate magnification serial block face SEM of a PtK_2_ cell. Black arrows highlight cytoskeletal elements extending into thin regions of the cell. White arrowheads = quantifoil carbon film, black asterisks = holes in the carbon film, M = mitochondria, ER = endoplasmic reticulum. C’ and C’’) High magnification transmission electron micrographs revealing the extended basolateral cytoskeletal elements to the cell periphery. Black arrows highlight cytoskeletal elements extending into thin regions of the cell. White arrowheads = quantifoil carbon film. D) Surface rendering of PtK_2_ cells. Black arrowheads = membrane extending over holes with thin cytoplasm; dark purple = no electron transparency; light purple = electron transparency. Scale = 2 µm.

We sought to optimise and maximise the number of compatible regions of interest for *in situ* cryo-ET without cryo-FIB lamella preparation. As the mechanical properties of cell substrates can be tuned to optimise important parameters such as cell spreading and cell-substrate adhesion, we first characterised the compliance/stiffness of the carbon film substrate on different regions of the carbon film across the grid square by atomic force microscopy (Fig 3A; Regions 1-3). We observed an increase in substrate stiffness to the carbon film overlaying the gold grid bar (Region 2) and the gold grid bar itself (Region 3) compared to the Young’s modulus of the carbon film in the centre of the grid square (Fig 3B, D, F and H; Region 1). This result aligns with our previous observations that mature focal adhesions preferentially form on the gold grid bar supported areas^9^ and observations made by others that cells preferentially align to the grid bar without modification of the substrate^10^. We next explored how changes in carbon substrate thickness could modulate support film stiffness properties. We deposited 12.5 nm of amorphous carbon onto a R2/2 gold support carbon film Quantifoil grid (heretofore after referred to as +Carbon). This addition resulted in an approximate ∼10-fold increase in the Young’s Modulus to the carbon film in the centre of the grid square (Region 1) compared to control grids with no additional carbon deposition (Fig 3B, C, I). We observed a two-fold increase in Young’s modulus on the grid bar (Region 2) comparing +Carbon and control grids (Fig 3D, E, J) and a reduction in stiffness on the gold grid bar (Region 3; Fig 3F, G, K). These data demonstrate that the thickness of the carbon substrate represents a mechanically tuneable property to control substrate stiffness to potentially manipulate cell adhesion properties.

**Figure 3.**
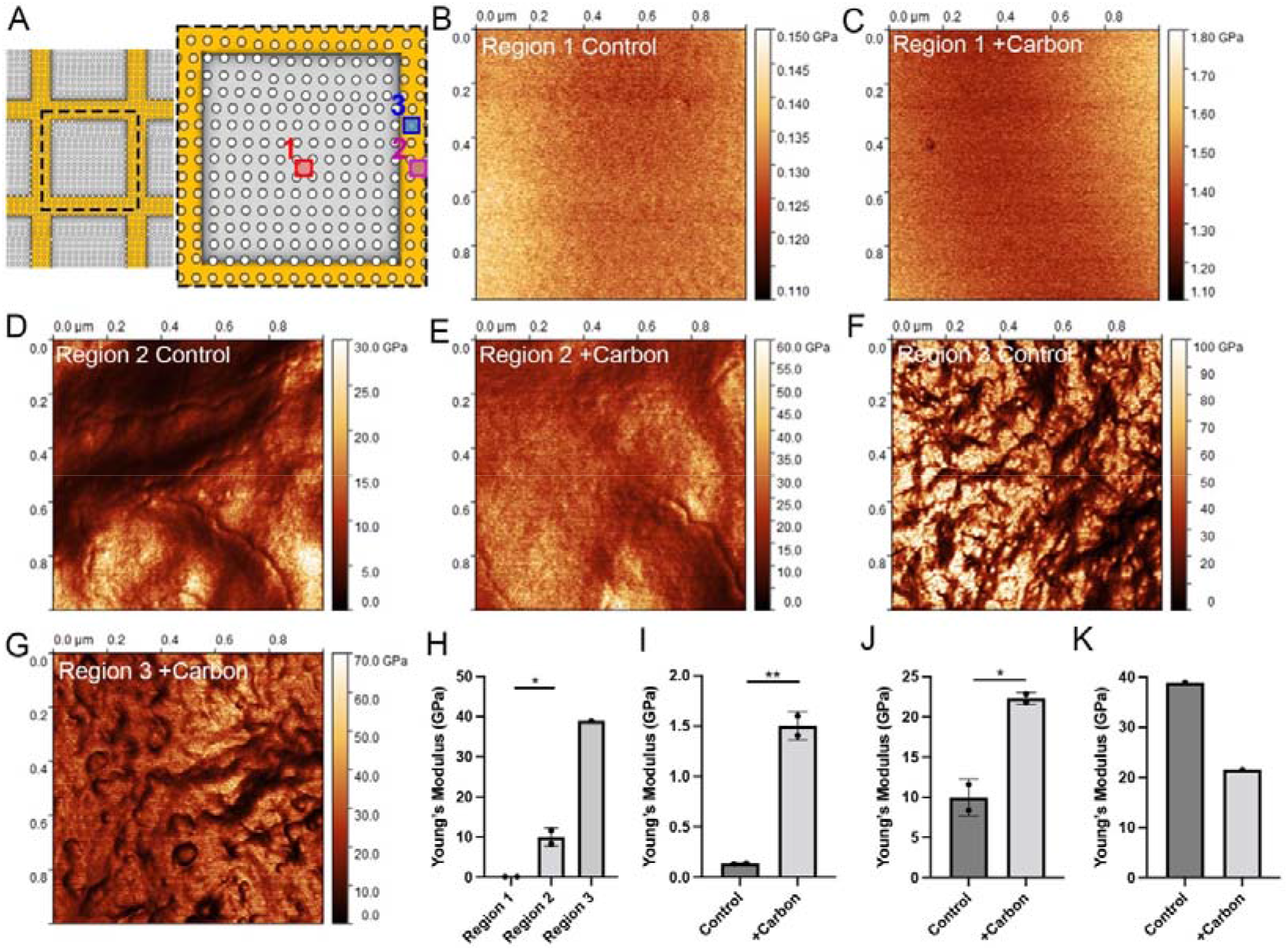
AFM of R2/2 AU 200 mesh carbon film Quantifoil grids. A) Schematic of Regions 1, 2 and 3 measured by atomic force microscopy. AFM measurements of a 1 µm^2^ area from B) Region 1 control, C) Region 1 +Carbon, D) Region 2 control, E) Region 2 +Carbon, F) Region 3 control and G) Region 3 +Carbon. H) Quantification of comparative changes in Young’s Modulus between Region 1, 2 and 3 from control grids. I-K) Quantification of comparative changes in Young’s Modulus between control and +Carbon from Region 1 (I), 2 (J) and 3 (K).

Given carbon addition modifies substrate stiffness, and substrate compliance is an important parameter to control cellular adhesion^11^, we next explored if additional carbon deposition could alter cellular distribution across the grid square. We mapped the spatial abundance of the centre of mass of nuclei of cells seeded onto control and +Carbon grids (see Methods; Fig S1A, B). Control grids demonstrated a preference for nuclei positioning at the edge of the grid square (n=209; Fig S1C, E). This preference was lost in cells grown on the stiffened +Carbon substrate (n=189; Fig S1D, E). Cells grown on stiffer substrates achieve greater cell spread and 2D surface area compared to those seeded on more compliant substrates^12^. This suggests with greater substrate stiffness we could increase the total number of areas suitable for cryo-ET by simultaneously reducing the number of cells localised to the edge of the grid square (where the ice is thickest) and increasing the spread of cells to reduce their thickness at the cell periphery. Substrate compliance is known to organise and remodel the actin cytoskeleton^13-15^. We therefore assayed stress fibre formation and actin organisation in PtK_2_ cells grown on control grids compared to +Carbon grids. Cells grown on control grids, and localised only to the carbon film with no association with the gold grid bar, demonstrated a significant reduction in total in phalloidin intensity per cell compared to similarly localised cells grown on +Carbon grids (Fig 4A-C). We also observed an increase in the proportion of stress fibres relative to whole cell area (Fig 4A, B and D) in +Carbon grids compared to those grown on control substrates. Strikingly, cells grown on control substrates that are localised to the gold grid bars demonstrated increased stress fibre formation compared to cells localised only to the grid square film (Fig 4A, C and D); this difference was less pronounced in cells grown on the stiffened substrate and localised to the gold grid bar (Fig 4B, C and D). These observations led us to investigate focal adhesion formation between cells grown on control and +Carbon grids. We assayed focal adhesion size with immunofluorescence against paxillin, which is a marker of focal adhesions and regulator of focal adhesion formation and stability. We observed a significant increase in the size of focal adhesions on the carbon film in cells grown on +Carbon grids compared to control Quantifoil grids (Fig 4E, F and G) suggesting that the additional carbon coat supports focal adhesion maturation. We also observed that focal adhesion size was rescued in cells grown on the control grids where the focal adhesions were localised to the gold grid bar (Fig 4H); this rescue was less pronounced when comparing between film and grid bar on +Carbon grids (Fig 4I). Therefore, we analysed the 2D spread area of cells grown on control and +Carbon stiffened substrates and observed a significant increase in 2D area to cells seeded on +Carbon grids (Fig 4J). Together, these data suggest that substrate stiffness is an important consideration for growing cells on grids and represents a tuneable parameter that can be used to modify cell size, cell spatial distribution over the grid and cell substrate adhesion. These optimisation steps aid data throughput by simultaneously increasing the number of thin peripheral areas suitable for cryo-ET analyses while bypassing the significant bottleneck of cryo-lamella preparation.

**Figure 4.**
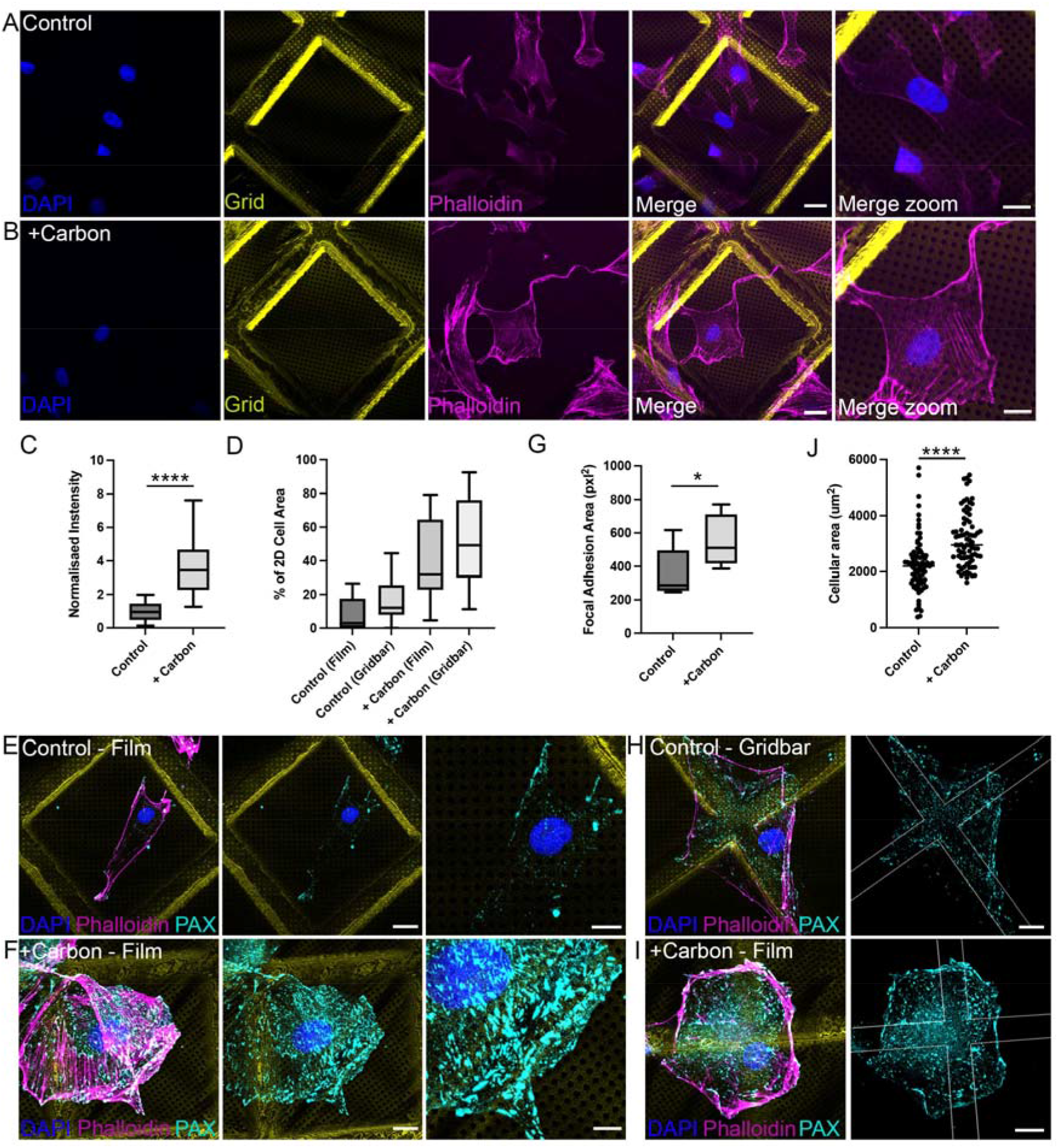
Substrate stiffness is a critical factor in regulating focal adhesions. A) Maximum intensity z-projection of a laser scanning confocal z-stack of cells seeded onto the control substrate demonstrating low phalloidin intensity (magenta) and poor stress fibre formation in cells growing exclusively on the carbon film. DAPI = blue; grid bar = yellow; phalloidin = magenta. Scale = 20 µm left panels; Scale = 10 µm in zoom. B) Maximum intensity z-projection of a confocal z-stack of cells seeded onto a +Carbon substrate. Film localised cells show greater phalloidin intensity and increased stress fibre formation compared to control cells. DAPI = blue; grid bar = yellow; phalloidin = magenta. Scale = 20 µm left panels; Scale = 10 µm in zoom. C) Quantification of whole cell background subtracted phalloidin intensity between control and +Carbon grids showing a significant increase in phalloidin intensity to cells grown on +Carbon. Statistical significance determined by two-tailed t-Tests. D) Percentage of cellular 2D area as stress fibre comparing control (film), control (grid bar), +Carbon (film) and +Carbon (grid bar) areas. E) Confocal z-stack of cells grown on Control grids showing minimal mature focal adhesions to the film. Scale = 20 µm left panels; Scale = 10 µm in zoom. F) +Carbon grids demonstrating increased immunofluorescence staining of paxillin. Scale = 20 µm left panels; Scale = 10 µm in zoom. G) Two-tailed *t*-test of focal adhesion size comparing control to +Carbon grids showing increased focal adhesion maturity with increased substrate stiffness. α-Paxillin = cyan; DAPI = blue; grid bar = yellow; phalloidin = magenta. H) Confocal z-stack of focal adhesions of cells grown on control grids show increased focal adhesion formation on the grid bar. Scale = 20 µm. I) Confocal z-stack of focal adhesions from +Carbon cells showing similar focal adhesion maturity between film and grid bar. Scale = 20 µm. J) Quantification of 2D cellular area comparing control grids to +Carbon grids.

### Rapid Cryo-CLEM

Having optimised cell-substrate interactions for improved yield of thin peripheral regions of interest for cryo-ET, we next pursued improvements in the speed and accuracy for imaging fluorescently labelled organelles of interest by cryo-CLEM (schematically depicted in Fig 1A and B). To test our rapid workflow, we pursued a data acquisition strategy that would encompass cryo-preservation to high-resolution data acquisition in less than a day. Cells were seeded onto +Carbon grids at low density to minimise potential cellular overlap. We observed that between 50-100 cells seeded per grid allowed for good quality and reproducible back sided blotting with sufficient areas for cryo-tomography (Fig S2A). For best imaging cells should be seeded below confluence. PtK_2_ cells were then transfected with EYFP-Mito7 as a marker of mitochondria, and 24 hours later and plunge frozen 24 hours post-transfection. Figure S2A demonstrates low magnification cryo-fLM image of EYFP-Mito7 transfected grid. Areas with thick ice are readily discernible by the high level of autofluorescence (Fig S2A). The grid was mapped, stitched, and regions of interest (ROIs) were selected for by low expression of EYFP-Mito7 with thin cell peripheries extending toward the centre of the grid square. Confocal z-stacks were acquired at ROIs 1 and 2 using a 100X objective (0.7NA 4mm long working distance) with the Airyscan 2 detector. This magnification with a digital zoom of 0.5 was sufficient to acquire whole grid square maps at a resolution high enough for targeting mitochondria (Movie S2; Movie S3). For high-resolution re-registration, both fluorescence and transmitted light channels must be acquired. This strategy allows for accurate correlation of the holes in the carbon film between cryo-fLM and cryo-EM imaging modalities and thus provides a sub-2µm fit corresponding to the diameter of the hole for all fluorescently labelled structures. Following this, the frozen grid was loaded into the cryo-TEM, a low magnification whole grid atlas was acquired (Fig S2B) and whole grid square ROIs were identified and re-registered (Fig S2A and B). Broken grid squares (white asterisks) represent a simple and rapid fiducial marker to correlate cryo-fLM images to cryo-TEM micrographs at the whole grid scale. 700X magnification cryo-TEM images were acquired at the ROIs and fitted to the grid square confocal z-stack using a combination of cellular landmarks and the position of the Quantifoil carbon film holes as correlation markers (Fig S2C and D). Images were fitted in real-time using Adobe Photoshop with Puppet Warping. From this data electron transparent regions in the cell periphery can be identified and correlated to EYFP-Mito7 signal for targeted analysis at the individual hole scale (Fig S2C’ and D’). ROIs 1 and 2 were suitable for cryo-tilt series acquisition (Fig S2A-D’; green and magenta squares), however ROIs 3 and 4 (Fig S2A and B; red and blue squares) represent areas too thick for cryo-ET; this was observed in the by z-stack (Movie S4, Movie S5 and Movie S6) and this was confirmed through 700X magnification correlation at the grid square level (Fig S2I-J’). For expedited analyses, ROI thickness should be screened and excluded during the cryo-fLM imaging. Fig S2E and F demonstrate intermediate magnification (2600X) overlay of ROI 2 showing EYFP-Mito7 signal correlating with electron dense structures with a mitochondrial morphology. From this overlay, Holes of Interest (HOIs) were selected for batch cryo-tilt series acquisition (Fig 7D and E). Batch tilts series were acquired overnight at each HOI with 20-30 tilt series were acquired per grid. HOIs 1 and 2 demonstrate high-magnification precision targeting of fluorescent signal and correlation with mitochondria (Fig S2G-H’’ respectively). This image acquisition strategy results in a highly accurate demarcation of larger organelles of interest quickly. For these assays, we plunge froze live cells in the morning and acquired batch tilt series overnight starting on the same day.

In the extended peripheral regions of the PtK_2_ cell cytoplasm, mitochondria occupy most of the cytoplasmic depth (Fig 1A and B), which effectively results in a near 2D correlation method. To extend our studies to a cellular structure that does not occupy the entire cytoplasmic volume, and thus an experimental condition that represents a more complex 3D re-registration process on a smaller structure, we performed cryo-CLEM analyses on GFP-tagged Doublecortin X (EGFP-DCX) as a marker of microtubules^16^. Using this method, we observed microtubules in our fluorescence images and correlated those signals to individual microtubules in the 3D volume of the reconstructed tomograms (Fig S3A–H; Movie S7). These data were also acquired in under a day and highlight that this protocol represents is robust and can be utilised to target smaller macromolecular complexes in the condensed, but still 3-dimensional, volume of the PtK_2_ cytoplasm.

### Optogenetic Positioning of Organelles-Cryo-CLEM (OPO-cryoCLEM)

The major constraint to the method described thus far is that imaging is restricted to the cell periphery. This places a physical limitation on our ability to image structures contained within thicker regions of the cell such as endolysosomes and recycling endosomes (Fig 1A). To enhance the utility of this novel PtK_2_ cell system we expanded our analyses to include perinuclear localised organelles without cryo-FIB. To do so, we sought to re-distribute perinuclear-localised organelles to the peripheral cytoplasm with optogenetic positioning of organelles (OPO). Commonly used systems to spatially modulate protein localisation include electro-genetic^17^, chemical genetic^18^ and photosensitive optogenetic switches^19^; the latter with the best spatial and temporal control (see^20^ for review). Optogenetic switches consist of a photoreceptor that undergoes a conformational change with stimulation of a particular wavelength of light, leading to interaction with specific partners via homo-dimerisation or heterodimerisation with bait and prey tags. Well-characterised optogenetic systems include CRY2/CIBN, iLID, and enhanced Magnets^21-23^. We have previously utilised the CRY2 (prey)/CIBN (bait) system to modulate endocytic recycling of the T cell receptor after T cell activation^24^. When coupled with bait-labelled kinesin motors, it is possible to redistribute prey-labelled organelles to the plasma membrane with inducible anterograde transport^21,25^. We observed abundant cytoskeletal elements extending all the way to the cell periphery (Fig 2C-C’’) and reasoned that we could utilise the CRY2/CIBN system coupled with anterograde motor transport to redistribute Lysosomal Associated Membrane Protein 1 (LAMP1)-labelled endolysosomes to the cell periphery using the bait-labelled kinesin motor, KIF5a (schematically depicted in Fig 5A). We first characterised LAMP1 redistribution with LAMP1-mCherry-CRY2 with KIF5a-GFP-CIBN expression in live and in fixed cells. We observed extensive anterograde transport of LAMP1 to the plasma membrane following light stimulation in cells by fluorescence microscopy (Fig 5B-E; Movie S8). Quantitation revealed a significant proportional redistribution of LAMP1 signal to the peripheral areas of the cell cytoplasm after 3 minutes of blue light stimulation (Fig 5F). We then confirmed that LAMP1-mCherry-CRY2 was redistributed to the cell perimeter with stimulation of blue light prior to plunge freezing with imaging by cryo-fLM and Airyscan 2 detection (Fig 5G and H). As an additional control we assessed HRP-labelled endolysosomal redistribution using conventional CLEM after HRP uptake and light stimulation. PtK_2_ cells transfected with both LAMP1-mCherry-CRY2 and KIF5a-GFP-CIBN were selected, subjected to a 30-minute HRP uptake at 37°C prior and 30-minute chase at 37°C to ensure HRP incorporation into LAMP1 positive structures. Cells were fixed after 3 minutes of light stimulation and cells of interest re-registered as performed previously ^26,27^. Unstimulated doubly transfected cells showed limited clustering of perinuclear HRP-labelled endolysosomes with minimal association with cytoskeletal elements (Fig 6A-C). Stimulated cells showed a dramatic association between clustered, HRP-labelled endolysosomes and cytoskeletal elements, as well as HRP-decorated vesicles highly enriched at the plasma membrane (Fig 6D-F). These data confirms that CRY2/CIBN can be used to reposition LAMP1 positive organelles to the cell periphery. We also confirmed this method could be used to redistribute Rab11-positive compartments (Fig S4; Movie S9).

**Figure 5.**
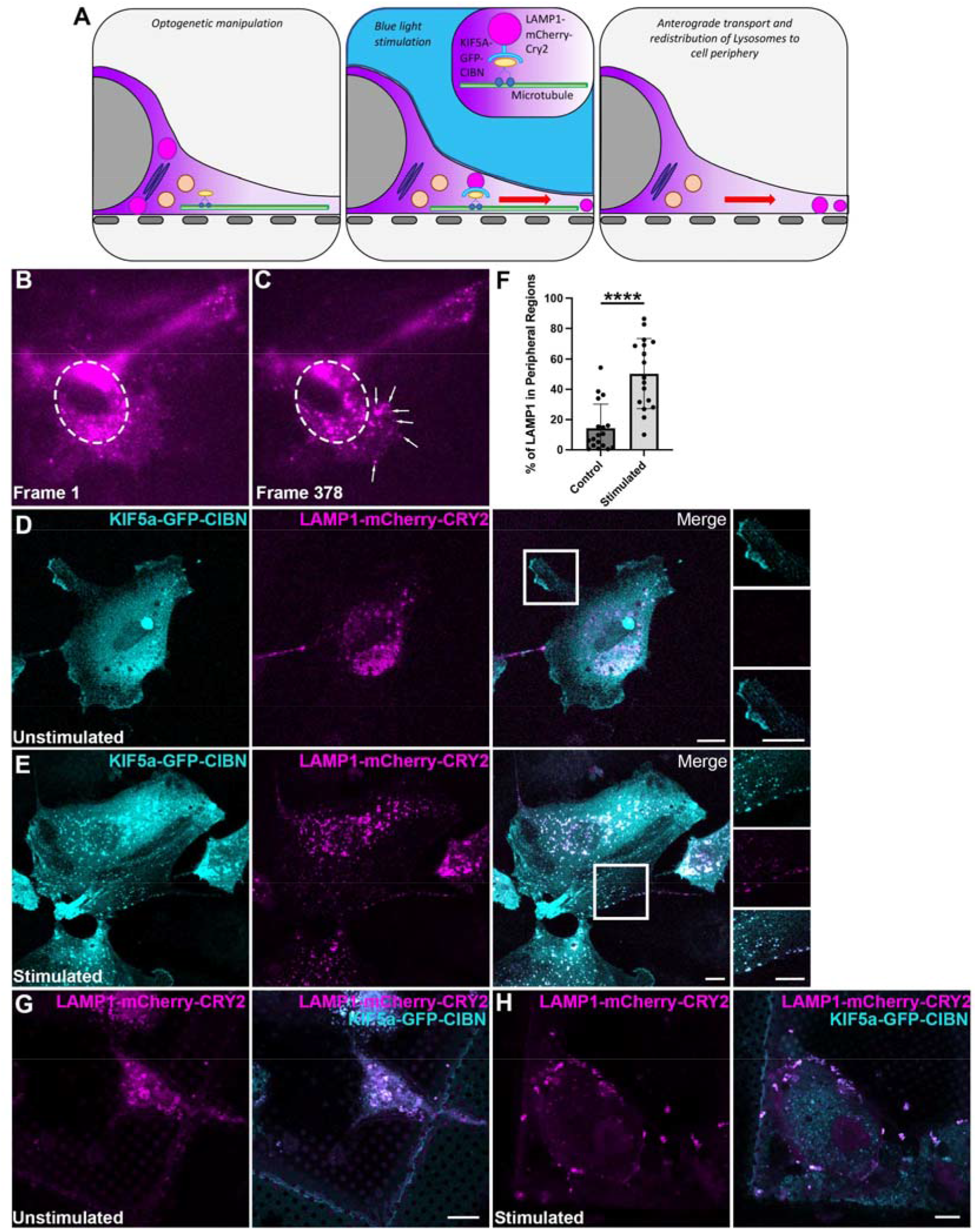
Optogenetic positioning of LAMP1 after light stimulation. A) Optogenetic positioning of endolysosomes from perinuclear regions to the cell perimeter. B) Live cell fluorescence image (frame 0) of a PtK_2_ cell expressing KIF5a-GFP-CIBN and LAMP1-mCherry-CRY2 immediately prior to optogenetic blue light stimulation. Dotted ellipse = perinuclear region. C) The same cell from (B) after 3 minutes (frame 378) of stimulation with blue light showing redistribution of LAMP1-mCherry-CRY2 from perinuclear regions to the cell periphery. Dotted ellipse = perinuclear region, White arrows = redistributed LAMP1 positive structures. D) Confocal microscopy of fixed PtK_2_ cells without unstimulated. LAMP1-mCherry-CRY2 is restricted to perinuclear regions. E) Confocal microscopy of fixed cells after 3 minutes blue light stimulation showing dramatic redistribution of LAMP-mCherry-CRY2 to the cell periphery. F) Quantification of LAMP1 fluorescence intensity comparing the percentage of peripheral signal to perinuclear signal. G) Cryo-confocal of plunge frozen grids seeded onto Quantifoil grids showing re-distribution of LAMP1-mCherry-CRY2 to the cell periphery blue light stimulation.

**Figure 6.**
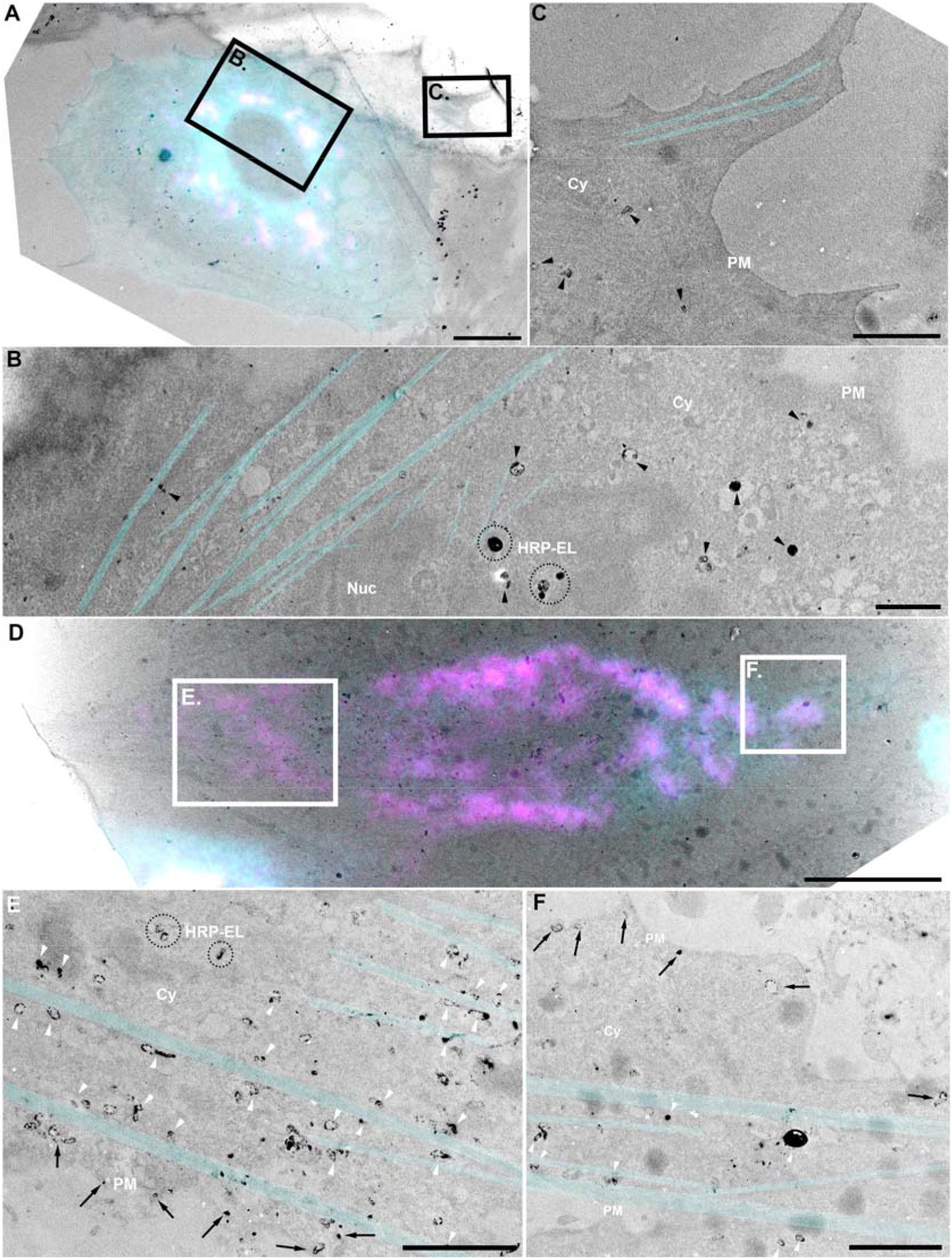
Optogenetic positioning of HRP-labelled endolysomes loaded with HRP. A) Conventional CLEM overlay of an unstimulated PtK_2_ cell expressing KIF5a-GFP-CIBN and LAMP1-mCherry-CRY2. Pulse-chase uptake showing HRP loaded lysosomes are located to perinuclear regions. Scale = 10 µm. White boxes = areas of interest. B, C) Higher magnification from (A) views showing HRP-positive structures are localised to the perinuclear region of the cell and are minimally associated with the cytoskeleton. PM = plasma membrane; Cy = cytoplasm; Nuc = Nucleus; HRP-EL = HRP-labelled endolysosomes; cyan = cytoskeleton; black arrowheads = HRP-labelled endolysosomes located to perinuclear areas. Scale = 2 µm. D) Conventional CLEM overlay of a PtK_2_ cell expressing KIF5a-GFP-CIBN and LAMP1-mCherry-CRY2 and stimulated with blue light for 3 minutes prior to fixation. Pulse-chase uptake showing HRP loaded lysosomes are tightly associated with microtubules and are located to the cell periphery. Scale = 10 µm. White boxes = areas of interest. E, F) Higher magnification views of white boxes from (D) showing association between lysosomes and cytoskeletal elements. PM = plasma membrane; Cy = cytoplasm; Nuc = Nucleus; HRP-EL = HRP-labelled endolysosomes; cyan = cytoskeleton; white arrowheads = microtubule-associated endolysosomes in the process of re-distributing to the cell periphery; black arrows = HRP-decorated peripheral endolysosomes. Scale = 2 µm.

Finally, we performed OPO-cryoCLEM and analysed the morphology of re-positioned LAMP1 positive organelles in doubly transfected cells. We observed LAMP1 in multiple locations at the cell periphery (Fig 7A-B). 3 ROIs were selected for batch cryo-ET acquisition (Fig 7C) and re-registered with our cryo-CLEM methodology; each LAMP1-mCherry-CRY2 structure localised at the cell periphery correlated with an electron dense vesicular structure at low magnification (Fig 7D and E). Cryo-tilt series were acquired at ROIs 1-3, reconstructed and overlayed with the cryo-fLM z-stacks. Endolysosomes were readily discernible at each mCherry-labelled vesicle (Fig 7F-H’; Movie S10-12) and these morphologies were consistent with other recent studies looking at lysosomal structure by cryo-FIB and cryo-ET^28,29^. To further demonstrate the utility of this method, we redistributed Rab11a-positive recycling endosomes from perinuclear regions to the cell periphery by co-expression of FuRed-Rab11a-CIBN, Kif5a-GFP-CIBN and Cry2-cluster-mCerulean. As with LAMP1, light stimulation resulted in significant redistribution of Rab11a to the cell perimeter, highlighting that this OPO-cryoCLEM is applicable to a diverse set of organelles. These data demonstrate that OPO-cryoCLEM can be used to re-distribute perinuclear localised organelles to the cell periphery in PtK_2_ cells, facilitating cryo-ET without the requirement for cryo-FIB and cryo-lamella preparation.

**Figure 7.**
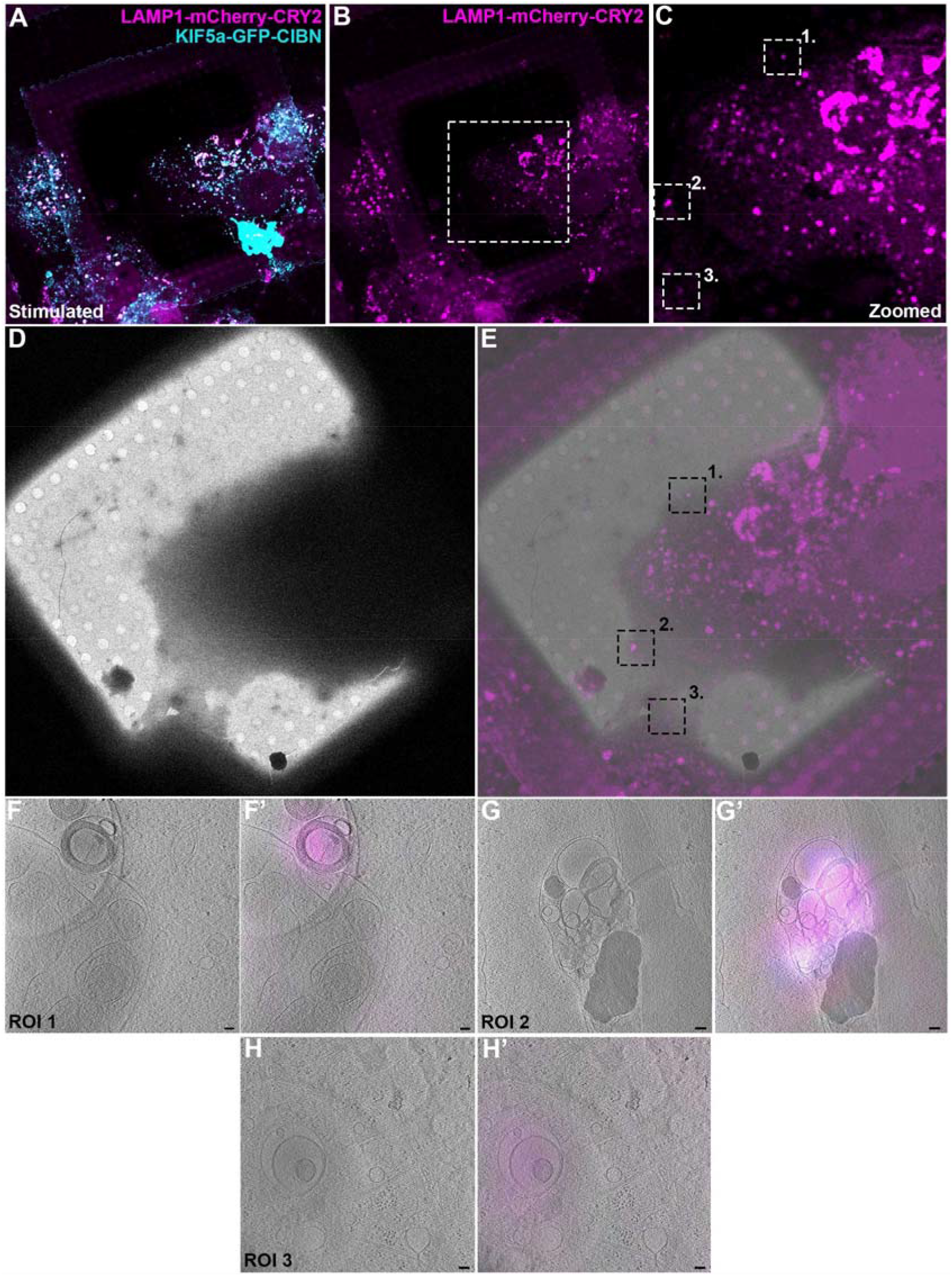
Optogenetic positioning of perinuclear localised organelles for cryo-ET without Cryo-FIB lamella preparation. A) Cryo-fLM of a blue light stimulated PtK_2_ cell expressing LAMP1 and KIF5a. B and C) LAMP1-mCherry-CRY2 signal showing 3 lysosomes localised to the cell perimeter (white dotted boxes). D) Cryo-electron micrograph of the cell from (A) at 700X magnification. E) Overlay of the LAMP1-mCherry-CRY2 signal with the cryo-EM image highlighting ROIs 1, 2 and 3 (Black dotted boxes). Optical slices of reconstructed tomograms showing the same regions of interest corresponding to ROI 1 (Hand H’), ROI 2 (G and G’) and ROI 3 (H and H’). Scale = 10 nm. The LAMP1 signal correlates with structures with a morphology consistent with endolysosomes.

## Discussion

*In situ* cryo-ET provides the cellular context to structural analyses that is lacking in traditional structural biology-based methods. These assays represent the next step towards the full exploration of the cellular environment at near atomic scale. To date, the field has been limited in its exploration of the cellular cryo-environment by a combination of factors including techniques with restricted throughput, hardware constraints and the expensive and challenging operational requirements of interconnected workflows between multiple high-end microscopes operated at cryogenic temperatures. Here we have developed a simple and novel PtK_2_ cell-based system that facilitates structural *in situ* analyses, from live cells through to batch cryo-tilt series data acquisitions, within 24-36 hours. By coupling PtK_2_ cells with optogenetic organelle positioning, we show that this system is not just restricted to *in situ* analysis of peripheral structures but can be used for perinuclear organelles like endolysosomes. The application of this system will allow for higher throughput examinations of diverse cellular structures and macromolecular complexes *in situ* and can be deployed at a fraction of the cost of the current standard in situ workflow.

This method has the advantage of side-stepping some of the most significant drawbacks associated with current *in situ* cryo-ET studies. First, the use of cryo-FIB for thinning samples which represents the largest time bottleneck for the wet lab component of the cryo-ET workflow^3^. Here, we developed a method that avoids the need for cryo-lamella preparation as an intermediate step by utilising the ∼200 nm thickness of the periphery of a PtK_2_ cell. Developing a protocol that avoids cryo-FIB has the flow on effect of avoiding other limitations associated with the existing methods, including the attrition rate for brittle cryo-lamella, the problems associated with preferential ice contamination of processed cryo-lamella, and the need for additional manual handling steps between microscopy modalities^3,30^. The throughput of this system is further improved by exploiting cryo-correlation between the cryo-confocal and cryo-ET such that ROIs are specifically targeted, bypassing off-target imaging and image processing. Second, this method results in only minimal ice contamination of our grids (see Figure S2, S3 and 7). By accelerating the workflow such that all grid transfers are performed in under 12 hours (plunge freezer, autogrid clipping, cryo-confocal imaging and the cryo-TEM autogrid loading), we could achieve high quality grids using only liquid nitrogen transfers without the requirement for high/ultrahigh vacuum transfer systems. Finally, the current gold-standard *in situ* cryo-ET workflow requires numerous expensive and highly specialised pieces of equipment including, a plunge freezer, a cryo-transfer system, a cryo-fLM capable system (either as a standalone system or incorporated into a cryo-FIB), a cryo-FIB and a cryo-TEM as well as the highly skilled staff to operate each instrument. This workflow, however, can be implemented within a facility at low cost, especially if the cryo-fLM stage can be utilised on an existing upright widefield fluorescence/confocal microscope.

The thickness of the PtK_2_ cell cytoplasm provides multiple advantages for cryoEM analysis. They possess a protracted and thin cytoplasm that provides ample electron lucent areas for *in situ* structural analyses, a cytoplasm densely packed with organelles, and represent a cell type that is highly amenable to genetic modification and expression of fluorescently labelled proteins of interest. While we have characterised this PtK_2_ cell system as ideal line for *in situ* cryo-CLEM analyses, we believe this technique can be applied to other cell types with similar morphological properties. As an initial screen we showed that MDCK, HeLa, BHK and Caco-2 cells were too thick for high quality cryo-ET imaging. We have previously performed cryo-CLEM on focal adhesions in MEFs^9^ and while sufficiently thin at the cell periphery, MEFs did not possess the ideal extended “fried egg” like morphology and as such are only appropriate for analyses directly at the cell perimeter. U-2OS cells also demonstrated an ultrastructural organisation similar to PtK_2_ cells. We suspect other lines with similar cell thicknesses, for example primary lymphatic endothelial cells^31^, would prove an appropriate line for *in situ* Cryo-CLEM/OPO-Cryo-CLEM analyses if cultured at sub-confluency on grids.

Cells grown on gold carbon grids preferentially associate with the grid bar^10^. Photo-micropatterning was developed to circumvent preferential cell adhesion through selective printing of extracellular matrix into the centre of a grid square to direct cell growth in regions better suited imaging adherent cell by cryo-ET^10^. Here, we sought to provide a characterisation of the cell-grid interface to determine why cells preferentially associate with the grid bar. Our studies demonstrated regional differences in substratum compliance, with the centre of the grid square as the most compliant region of the film with the carbon film over the gold grid bar and the gold grid itself to be orders of magnitude greater in substrate stiffness (Fig 3). These stiffer areas also demonstrated an increased total number, and maturity of, focal adhesions, increased phalloidin intensity and greater stress fibre formation (Fig 4) this is commonly observed comparing between glass coverslip substrates and collagen matrices. This indicated that substrate stiffness may represent a tuneable parameter to modify how cells associate with the Quantifoil carbon substratum. As such we altered carbon film thickness through deposition of additional amorphous carbon and observed that evaporation of 12.5 nm thick carbon layer resulted in an approximately 10-fold increase in substrate stiffness that translated into increased focal adhesion formation, increased focal adhesion maturity, increased phalloidin intensity and increase in stress fibre formation. Additionally, these changes resulted in a redistribution of the centre of mass of the nucleus from the edge of the grid square for cells seeded on control grids toward a more random distribution in +Carbon grids and a commensurate increase in the 2D cell area. This indicates that additional carbon deposition may represent a low-cost option to increase the yield of suitable areas for cryo-ET. Our studies also show that additional carbon deposition can be used experimentally to control substrate stiffness allowing investigation of *in situ* structural changes in mechanosensitive and mechanoresponsive macromolecular assemblies.

In the OPO-Cryo-CLEM system we utilised optogenetics to position organelles in the periphery of the cell making them amenable to cryo-electron tomography. The system could conceivably be modified to redistribute and structurally characterize non-membranous structures or protein complexes. While several optogenetic systems have been used to stimulate redistribution of organelles and proteins of interest to specific subcellular compartments^22,23,32-38^, the temporal properties of the CRY2/CIBN system proved favourable for our purpose. The CRY2/CIBN interaction is transient and reversible, with interaction occurring within seconds and reversion taking minutes to hours^21^ compared to other systems which can reverse within seconds to minutes timescale^23^. For our purposes, the slower reversion/off rate of the CRY2/CIBN system ensured ample time between stimulation and plunge freezing to minimise redistribution of organelles back to the perinuclear space.

### Caveats

As with any new method, certain caveats must be considered before its application. Firstly, the choice of cell system is clearly very important. Thicker cells will not allow sufficient electron penetration without cryo-FIB milling. The size of the organelle of interest also needs to be carefully considered as thicker organelles will result in lower quality cryo-ET imaging. Here we focused on the redistribution of endolysosomes which are large organelles on the upper limit of what is possible. Larger lysosomes (above 300nm in diameter) may not be able to be imaged using cryoEM due to their thickness. To avoid electron transparency issues, we focused our analyses specifically on the smaller endolysosomes recruited to the cell periphery after optogenetic stimulation. This was less of an issue with OPO-cryoCLEM as the larger endolysosomes were not actively transported to perinuclear regions compared to the smaller endolysosomes (see Fig 5). For some applications the disruption of the steady state distribution of an organelle or protein complex during optogenetic repositioning could be disruptive, especially to transient assemblies and highly dynamic structures (for example sorting tubules). We did however observe significant re-distribution of HRP-loaded endolysosmes within 3 minutes of light stimulation suggesting that this technique is well-suited for analyses of cargo within a membrane-enclosed structure. Despite these caveats, the use of OPO-cryoCLEM has the potential to open a new era of structural characterization of a wide range of protein and membrane assemblies inside native cells.

## Methods

### Carbon coating

Carbon coating performed on a SafeMatic CCU-10 carbon coater fitted with a CT-010 Carbon Thread evaporation module. R 2/2 carbon film gold support Quantifoil grids (Q2100AR2; Electron Microscopy Sciences) were loaded into the vacuum chamber and carbon was deposited under high vacuum with stage rotation. Carbon deposition was automatically detected by the Quartz film monitor sensor and deposition was stopped with the thickness monitor reached 12.5 nm.

### Atomic Force Microscopy

Carbon and control Quantifoil grids were mounted on a PDMS substrate for analysis in a Bruker Dimension ICON SPM equipped with a Nanoscope V controller. The sample is allowed to settle on the PDMS substrate for at least 2 days to avoid drifting during AFM measurements. Surface topology and the DMTmodulus were acquired using Peakforce QNM mode, in 1 µm^2^ areas in duplicate at 3 regions of the grid square using OTESPA-R3 tips (from Bruker AFM tips); Region 1 in the centre of the grid square on the carbon film; Region 2 on the carbon film over the gold support bar; Region 3 in the Quantifoil hole over the gold grid bar (1 µm^2^ areas are represented in Fig 3A). Average stiffness measurements were generated from each point within the 1 µm^2^ area corresponding to 256 positions in X and Y. Significance was determined by two-tailed Student t-tests. The AFM images were processed with Gwyddion (version 2.59) software.

### Cell Culture

Low passage PtK_2_ cells (male potoroo kidney epithelial cells; ATCC) were cultured in Eagle’s Minimum Essential Medium supplemented with 10% Foetal Bovine Serum (Gibco) and MEM Vitamin Solution (Sigma Aldrich). Cells were trypsinized (0.05% Trypsin-EDTA; Sigma Aldrich) and seeded onto control and +Carbon grids as described in Figure 1B. Briefly, grids were placed in 35mm plastic tissue culture dish (Nunclon) and subjected to ultraviolet light for 20 minutes to ensure sterility. A 40 µL droplet of cells resuspended in MEM medium was placed over each grid with the carbon side facing up for 2-3 hours prior to the gentle addition of 2 ml of MEM. Cells were transfected with Lipofectamine 3000 (Invitrogen) as per the manufacturer’s instructions 24 hours after cell seeding. The plasmids used in this study were as follows: YFP-Mito-7 was a gift from Michael Davidson (Addgene plasmid # 56596 ; http://n2t.net/addgene:56596 ; RRID:Addgene_56596), EGFP-DCX was a gift from Joseph Gleeson (Addgene plasmid # 32852 ; http://n2t.net/addgene:32852 ; RRID:Addgene_32852), Kif5A-GFP-CIBN was a gift from Bianxiao Cui (Addgene plasmid # 102252 ; http://n2t.net/addgene:102252 ; RRID:Addgene_102252), LAMP1-mCherry-CRY2 was a gift from Bianxiao Cui (Addgene plasmid # 102249 ; http://n2t.net/addgene:102249 ; RRID:Addgene_102249). BHK, MEF, MDCK and Caco-2 cells were passaged as described above but grown in DMEM supplemented with 10% Foetal Bovine Serum (Gibco) and L-Glutamine (Invitrogen). U2OS were grown in McCoy’s 5a medium supplemented with 10% foetal bovine serum.

### Serial block face scanning electron microscopy

Sample preparation was performed as described previously^39^. Briefly, cells were seeded onto Quantifoil grids as described above, fixed in 2.5% glutaraldehyde in in phosphate buffered saline (PBS; pH 7.4), washed 3x in PBS and postfixed in 2% OsO_4_ with 1.5% potassium ferricyanide. Samples were then washed in double distilled H_2_O, incubated in a 1% (w/v) thiocarbohydrazide solution and postfixed again in 2% OsO_4_. Cells were washed (ddH_2_O) and stained *en bloc* with 1% uranyl acetate and subsequently with lead aspartate solution (20 mM lead nitrate, 30 mM aspartic acid, pH 5.5). Grids were serially dehydrated in ethanol, serially infiltrated with durcupan resin and polymerised for 48 hours. SBF-SEM imaging was performed as described previously^40^. Image stacks were assembled and aligned in IMOD. Segmentation was performed with Isosurface rendering in IMOD.

### Conventional TEM

Cells processed were processed as described above then Durcopan embedded blocks were subjected to ultramicrotomy. Vertical ultrathin sections (60 nm) were cut on a UC6 Leica Ultramicrotome and mounted on copper 200 mesh grids. Grids were imaged on a JEOL JEM-1011 Transmission Electron Microscope fitted with an EMSiS Morada camera.

### HRP uptake and conventional CLEM

Cells were seeded onto 35 mm gridded in plane MatTek dishes. Cells were transfected with LAMP1-mCherry-CRY2 and KIF5a-GFP-CIBN, left for 48 hours post-transfection, and then pulsed with 5 mg/mL of HRP in MEM media for 30 minutes at 37°C. The uptake media was replaced with normal MEM growth media and the cells were chased for a further 30 minutes at 37°C to ensure HRP incorporation into lysosomes. PtK_2_ cells were stimulated with blue light for 3 minutes (control cells were left unstimulated in MEM media) and fixed in 4% paraformaldehyde (PFA) with 0.1% glutaraldehyde in PBS for 30 minutes. Cells were washed in PBS and imaged on a Nikon Ti inverted fluorescence microscope. Cells positive by fluorescence for KIF5a and LAMP1 were imaged, and their alphanumeric code recorded by brightfield. Dishes were then processed using the DAB reaction as described previously^41^ and correlation was performed as described previously^27^.

### Fluorescence Microscopy

*Fixed Cells*. Cells were seeded on control or +Carbon grids as described above for Fig 4 and S1 and seeded onto coverslips for Fig 5. Cells were fixed with 4% PFA, washed in PBS, permeabilised with 0.1% Triton X-100 and stained with phalloidin (ABCAM) or DAPI (Thermo Fisher) as per the manufacturer’s instruction. For mounting, grids were inverted onto coverslips with mowoil and coverslips were then inverted onto slides such that cells were facing toward the coverslip and no fluorescence signal could be obscured by the gold grid bars. For nuclei positioning, mounted grids were imaged on a Nikon Eclipse Ti inverted fluorescence microscope. Grid squares were then divided into a 12 × 12 matrix of equal size (10 µm × 10 µm). The centre of mass of the nucleus was determined ImageJ and the coordinates of all CoMs were recorded and overlayed on the matrix. The 144 positions were rotationally averaged into 4 quadrants of 36 individual coordinates. These positions were then analysed by 5 groupings relative to the number of matrix positions away from the grid bar. Total phalloidin fluorescence intensity was normalized to background and analyzed per cell. *Immunofluorescence*. Cells grown on control or +Carbon were fixed in 4% PFA, washed in PBS, permeabilised in 0.5% Triton X-100, washed in PBS, quenched in 50 mM NH_4_Cl in PBS for 10 minutes and blocked in 1% bovine serum albumin in PBS. Grids were incubated with mouse α-Paxillin antibody (Thermo Fisher; Cat # AHO0492) at a dilution of 1:200 in the blocking solution for 30 minutes, washed in block and probed with a Goat α-mouse Alexa Fluor 647 conjugated secondary antibody (ABCAM; Cat # AB150079). Grids were PBS washed and stained and mounted as described above. Confocal z-stacks were acquired on a Nikon C2-Si upright confocal. *Optogenetics*. Stimulation was performed for 3 minutes with 405nm laser excitation on either an inverted Nikon Eclipse Ti with a mercury lamp or an Olympus CKX53 fitted with a CoolLED *p*E-300^white^ LED fluorescence module as described previously^21^. *Live cell*. PtK_2_ cells were plated on 13 mm coverslips and transfected with LAMP1-mCherry-CRY2 and KIF5A-GFP-CIBN or FuGeneRed-CIBN-Rab11a, KIF5A-GFP-CIBN and Cry2CIust-mCerulean using Lipofectamine 3000 overnight prior to imaging. Samples were transferred into fresh MEM containing 10% serum and MEM Vitamins and without phenol red 1 hour prior to imaging. Imaging was performed at 37°C. Simultaneous imaging and optogenetic stimulation was performed with HILO on a Zeiss ELYRA P1 microscope during widefield imaging with 200 mW 405 nm, 200 mW 488 nm and 200 mW 561 nm laser (LAMP1) or Zeiss 780 LSM with an Argon ion 488 nm and diode 561 nm laser. Samples were imaged for up to 1000 frames (ELYRA) or 100 frames (780 LSM). Samples were imaged for up to 1000 frames. Quantification of redistribution of endolysosomes was performed as described preiovusly^21^.

### Plunge Freezing

Grids were prepared as described above. Plunge freezing was performed on a Leica EM GP with back sided blotting. The chamber temperature was set to 37°C, humidity was 95% and blot time was varied between 4s and 6s with a 0s post-blot time. Grids were plunged into liquified ethane at -182 °C. Grids were transferred to liquid nitrogen for autogrid clipping and were clipped as per the manufacturer’s instructions (Thermo Fisher). Blue light stimulation prior to plunge freezing was performed on an Olympus CKX53 optical microscope fitted with a CoolLED *p*E-300^white^ LED fluorescence module. This optical microscope was set up beside the Leica EM GP. Blue light stimulation was performed for 2.5 minutes and grids were transferred onto the Leica EM GP for immediate freezing. Grids were cryo-preserved within 3 to 3.5 minutes of starting light stimulation.

### Cryo-Confocal imaging

Clipped autogrids were loaded in the Linkam CMS196 cryostage at -196°C and transferred onto the liquid nitrogen cooled bridge. The cryo-stage was mounted onto a Zeiss LSM 900 upright confocal fitted with 10X EC Plan-Neofluar (0.3NA; Zeiss) and 100X LD EC Epiplan-Neofluar (0.75NA; Zeiss) objectives and an Airyscan 2 detector. Whole grid mapping was performed as follows; the 10X objective at a 0.5 digital zoom was used to image the whole grid with T-PMT transmitted light and fluorescence channels as this objective and digital zoom allows for fast atlas acquisition with only 4 images required to image the whole grid. Focal points were set for each grid quadrant using Advanced setup in Zen Blue 3.2 and acquired with preview scan to generate an interactive map of coordinates, and subsequently the whole grid was acquired as a tile set and stitched in Zen Blue. ROIs/grid squares of interest were selected, and the z-stacks were acquired at each ROI using the 100X objective at 0.5 digital zoom with Airyscan detection of both fluorescence and T-PMT channels to image the whole grid square. Higher magnification z-stacks were acquired at a 1.6X digital zoom using the 100X objective with Airyscan and processed for Airyscan deconvolution. Excitation lasers were 488 nm (GFP/YFP) and 561nm (mCherry) with pixel dwell time 1.31 µs, and a z-slice optical section thickness between 100 nm and 500 nm. Pixel arrays of 1586 × 1586 pixels (125.88 µm × 125.88 µm) at 50X and 2024 × 2024 pixels (39.46 µm × 39.46 µm) at 160X. Up to 10 ROIs were imaged per grid. Z-stacks required between 30 minutes to 2 hours to acquire all ROIs per grid. We observed that ice contamination increased dramatically when the Linkam CMS196 cryo-stage was used with manually filling. The use of Linkam Autofill Dewar resulted in minimal ice contamination. Grids were transferred to liquid nitrogen for cryo-EM.

### Cryo-TEM, Cryo-ET, and real-time cryo-CLEM

Clipped autogrids were loaded into a 200□kV Talos Arctica Cryo-TEM microscope (Thermo Fisher). Whole grid batch atlases were acquired to relocate the grid squares of interest. Orientation and rotation were established using broken grid squares as fiducial markers. Atlases were exported to Adobe Photoshop and overlayed with the corresponding low magnification cryo-fLM map to establish grid squares of interest. Whole grid square ROIs were imaged using the Overview magnification at 700X in Tomography (Thermo Fisher) and this data was exported in real time to Adobe Photoshop for intermediate magnification alignment with maximum intensity z-projected cryo-fLM z-stacks. Correlation was performed using cellular landmarks and carbon film holes as markers in space. Aberrations between imaging modalities were corrected using Puppet Warping. The aligned z-projected fluorescence image overlayed on the 700X Overview cryo-EM image was sufficient to determine the positioning of fluorescently labelled structures and electron transparency: this manual alignment method allowed for rapid re-registration at sub-hole resolution to determine holes of interest. Batch cryo-tilt series were acquired with dose-symmetry in the Tomography software (Thermo Fisher). Single-axis tilts were collected from −60° to +60° at 2° increments at 7 □µm defocus on a Falcon 3 camera (Thermo Fisher) operated in linear mode at 22,000X magnification. Total electron dose was kept below 50 *e*^−^□Å^−2^ to minimise radiation damage. Tilt increments were aligned through a combination of fiducial tracking of 10 nm gold particles and Patch Tracking in etomo. Tilt series were reconstructed using weighted back-projection and filtered for visualization with SIRT-like filters and nonlinear anisotropic filtering in IMOD. Magnification scaling at the individual hole level was utilised to correlate cryo-fLM with the reconstructed cryo-tomogram.

## Acknowledgements

The authors acknowledge; the use of the Cryo-Electron Microscopy Facility through the Victor Chang Cardiac Research Institute Innovation Centre, funded by the NSW government, and the Electron Microscope Unit within the Mark Wainwright Analytical Centre (MWAC) at UNSW Sydney; the facilities, and the scientific and technical assistance, of Microscopy Australia and the Centre for Microscopy and Microanalysis, The University of Queensland, and; the Australian Cancer Research Foundation (ACRF)/Institute for Molecular Bioscience Cancer Biology Imaging Facility, which was established with the support of the ACRF. This work was supported by the National Health and Medical Research Council of Australia grants APP1185021 to NA; APP1140064 and APP1150083 and Fellowship APP1156489 to RGP). N.A. is supported by a UQ RS Fellowship. MLC was supported by a UNSW Scientia PhD Scholarship. P.W.G. and E.C.H. were supported by grants from the ARC (DP160101623), the NHMRC (APP1100202, APP1079866) and The Kid’s Cancer Project. VA is funded by the EMBL Australia Programme at UNSW.

## Contributions

N.A., R.G.P., G.M.I.R., and J.Rae conceived the study. N.A., J.Rae and G.M.I.R. performed cell culture. N.A. and G.M.I.R. performed light microscopy and V.A. aided with interpretation of fluorescence data. J.Rae and N.A. performed conventional electron microscopy. Y.Y. performed atomic force microscopy. N.A. and J.Ruan performed cryo-ET. N.A. and R.W. designed cryo-fLM acquisition strategy and N.A. performed cryo-confocal microscopy. N.A. and M.L.C. performed analysis of cell-substrate data and P.W.G. and E.C.H. aided with interpretation of substrate data. N.A. wrote the manuscript. N.A., R.G.P., V.A., P.W.G., and G.M.I.R. edited the manuscript.

## Supplemental Figures

**Figure S1.**
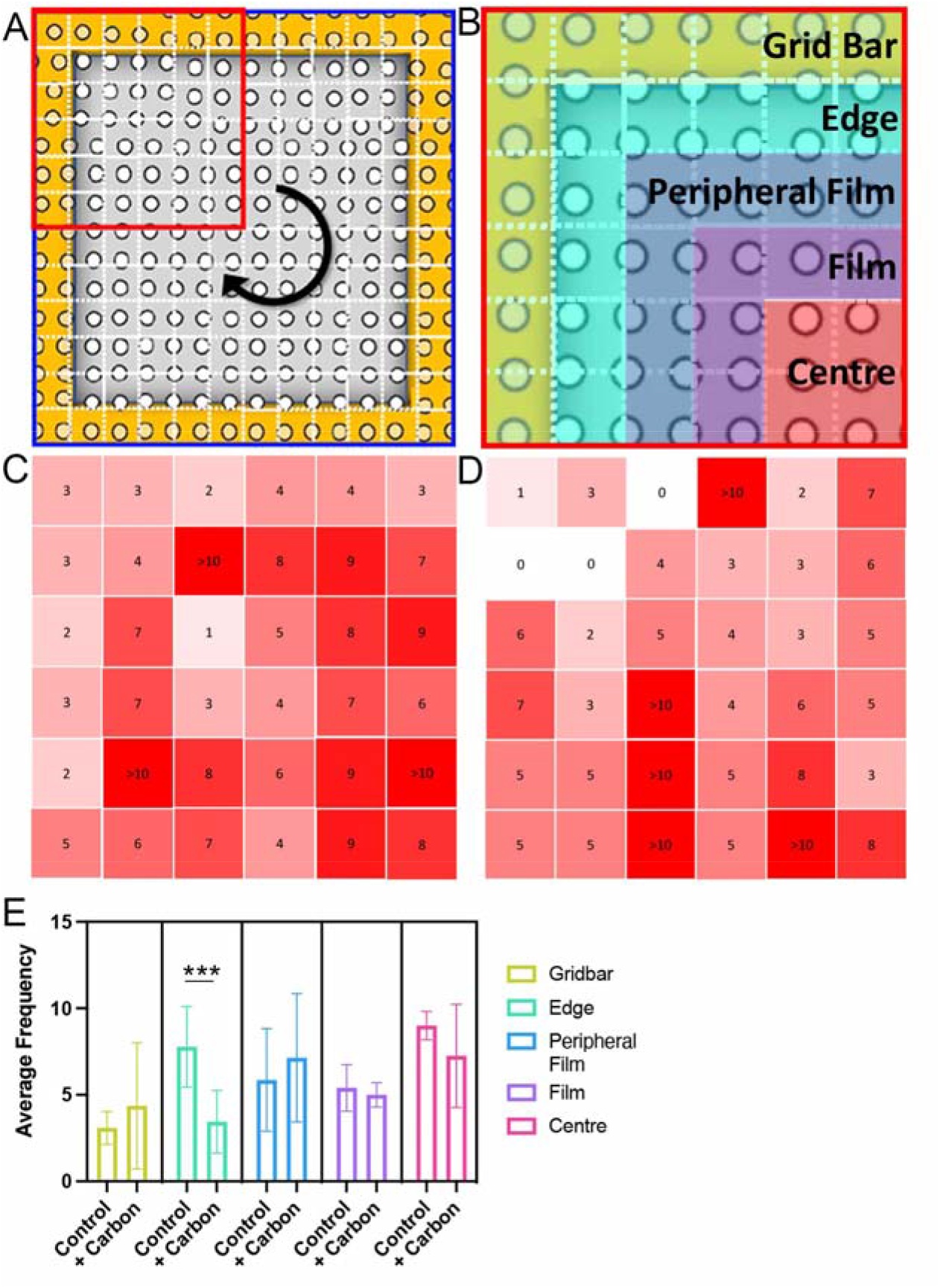
Substrate stiffness as a tuneable property to alter cellular distribution and spread on grid squares. A) Schematic of grid square divided into 144 coordinates (12 × 12 grid; white squares) of equal size (∼10 µm x ∼10µm) spanning the midpoint of the grid bar to the corresponding midpoints of the adjacent grid bars. Using the centre of mass of the nucleus as an approximation for the centroid of a cell, cell positions were mapped between control and +Carbon grids at equivalent confluence. The 144 coordinates were divided into 4 quadrants (red square) and rotationally summed to provide 36 final positions for scoring centre of mass. B) Positional classes based on relative distance from the grid bar. C) Heat map of frequency of centre of mass for each 36 positions for control grids. D) Heat map of frequency of centre of mass for each 36 positions for +Carbon grids. E) Quantification of relative abundance comparing cells from each position class. A significant reduction in “Edge” localised cells was observed in +Carbon grids.

**Figure S2.**
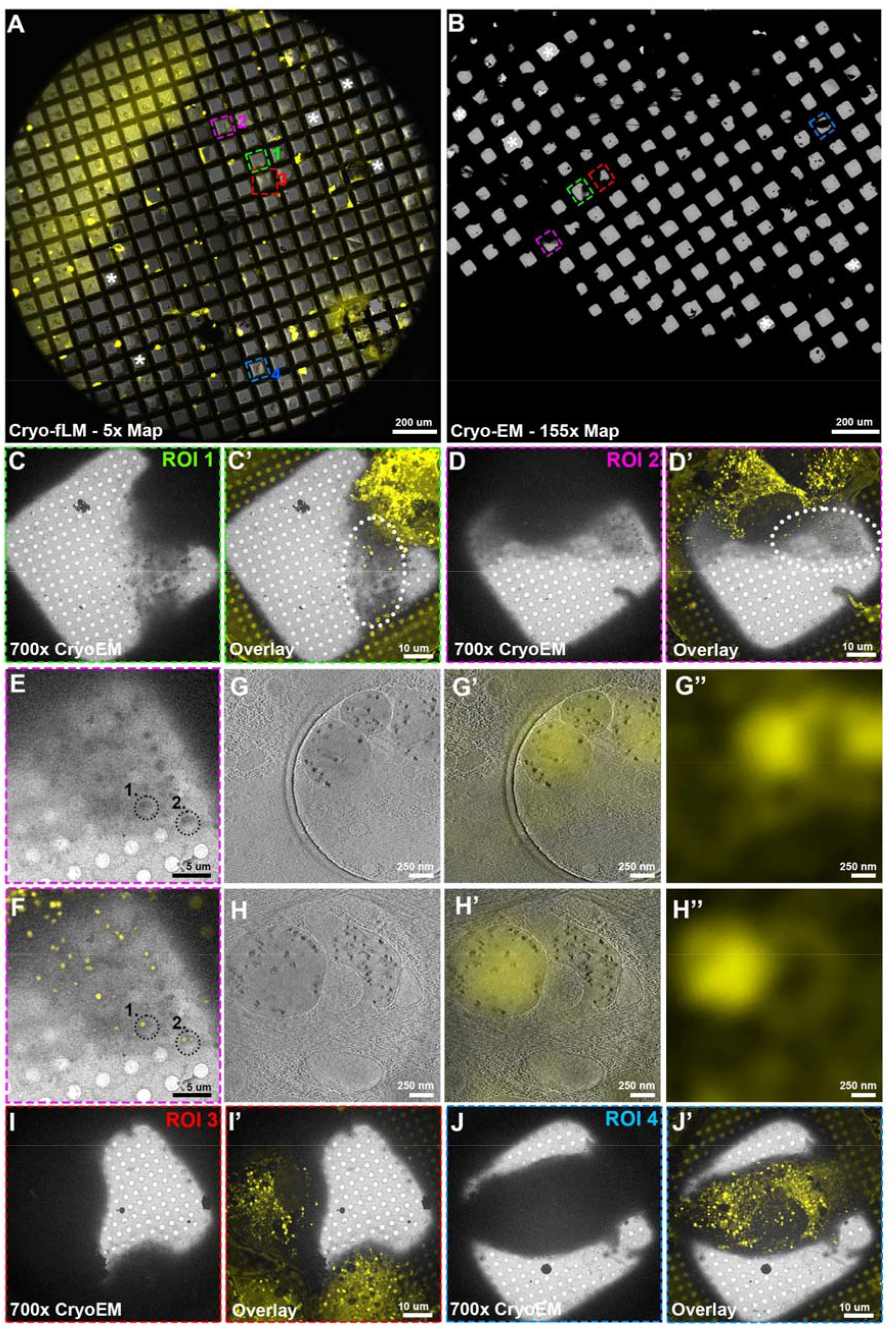
Low to high magnification Cryo-CLEM of EYFP-Mito7. A) Stitched confocal tile set of a whole grid of PtK2 cells expressing EYFP-Mito-7 grown on +Carbon film Quantifoil grids. Four regions of interest are highlighted representing transfected cells (ROI 1-4; green, magenta, red, blue, respectively). White asterisks demark holes for re-registration between the cryo-fLM and cryo-TEM. B) Stitched low magnification cryo-EM atlas of the same grid demonstrating the same four ROIs from (A). C, C’) Intermediate magnification of ROI 1 grid square showing the cryo-TEM and overlay of maximum intensity z-projections of confocal z-stacks. The electron transparent peripheral region enclosed within the white ellipse demonstrating EYFP-Mito-7 puncta. D, D’) Cryo-TEM and cryo-fLM overlay of ROI 2 demonstrating an electron lucent cellular periphery enclosed within the white ellipse area also positive for EYFP-Mito-7. E and F) Intermediate magnification Cryo-TEM (E) and cryo-fLM (F) overlay of the EYFP-Mito-7 expressing PtK_2_ cell from ROI 2 from (D,D’). Black dotted HOIs 1 & 2 are examples of suitable holes for batch cryo-tilt series acquisition. Scale = 5 µm. G) Optical slice from the reconstructed tomogram of the black dotted HOI 1 from (E) and (F). G’) High magnification overlay of EYFP-Mito7 signal with HOI 1 from the reconstructed tomogram in (G). G’’) Confocal signal from HOI 1. Scale = 250 nm. H) Optical slice from the reconstructed tomogram of the black dotted HOI 2 from (E) and (F). H’) High magnification overlay (H’) of EYFP-Mito7 signal with HOI 2 from the reconstructed tomogram in (H). H’’) Confocal signal from HOI 2. Scale = 250 nm. Thick cell peripheries need to be avoided for fast cryo-ET acquisition. I and J) Low magnification cryo-fLM and cryo-TEM maps as in A and B. Z-stacks should not be acquired where ROIs demonstrate a large deviation in z away from the plane of the carbon film (Movie S3 and 4). ROIs 3 (I and I’) and 4 (J and J’) correspond to cells that are too thick for cryo-electron tomography without cryo-FIB milling. Minimising thick cells expedites data acquisition.

**Figure S3.**
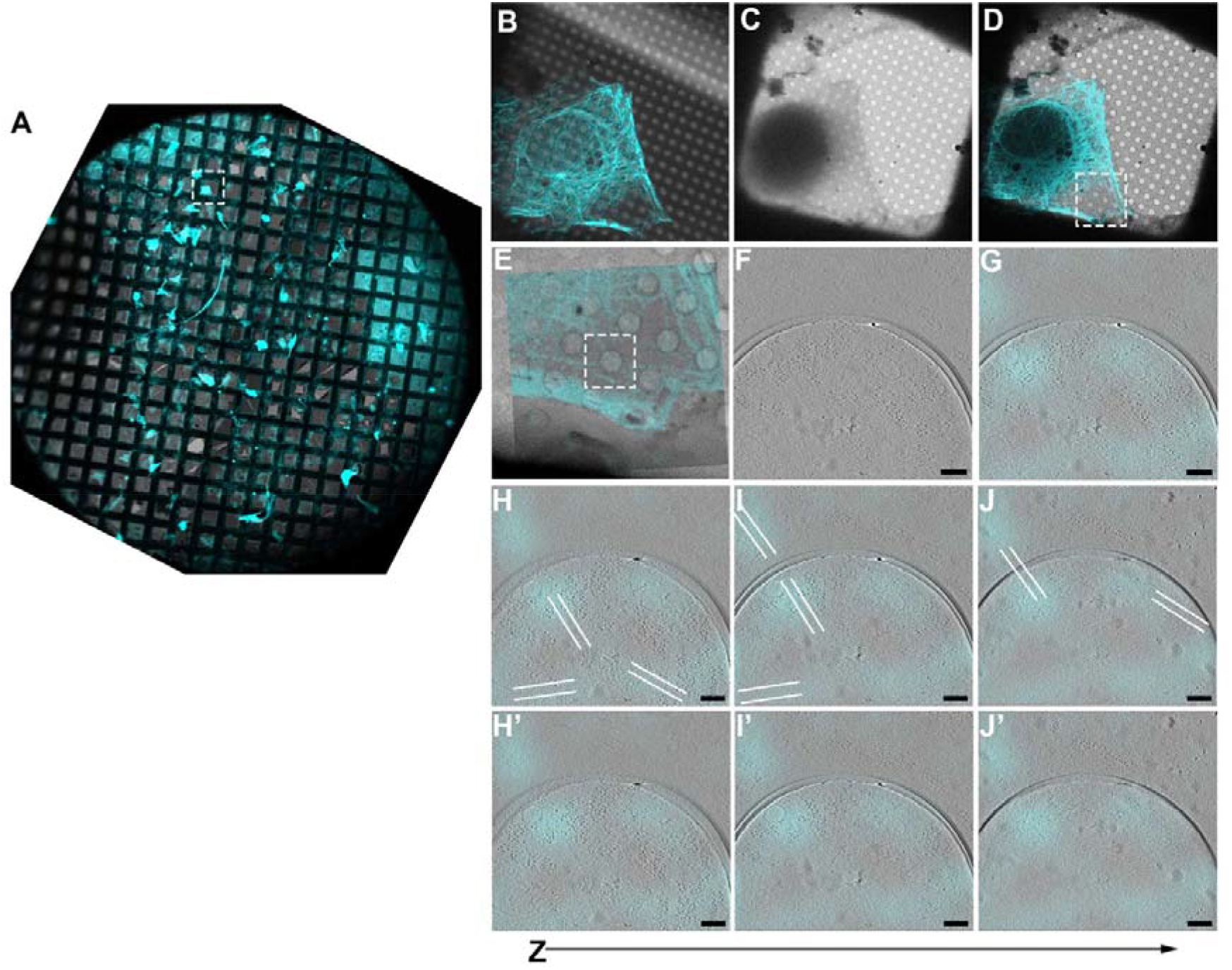
3D cryo-CLEM of microtubules. A) Whole grid stitched cryo-fLM map of PtK2 cells expressing EGFP-DCX. B) Confocal z-stack of EGFP-DCX signal with transmitted light demonstrating abundant microtubules. C) 700X magnification grid square view of the same cell from B). D) Overlay of B & C. White box = area of interest with thin cytoplasm and abundant microtubules. E) Intermediate magnification fitting at 2600X overlaying the z-projected confocal stack onto the search magnification image. White box = hole of interest. F) Optical slice of reconstructed tilt series acquired at the hole of interest from (E). Scale = 100nm. G-J’) Overlay of the EGFP-DCX signal with the reconstructed tomogram showing individual microtubules at the site of enriched fluorescence signal. White dotted lines = microtubules.

**Figure S4.**
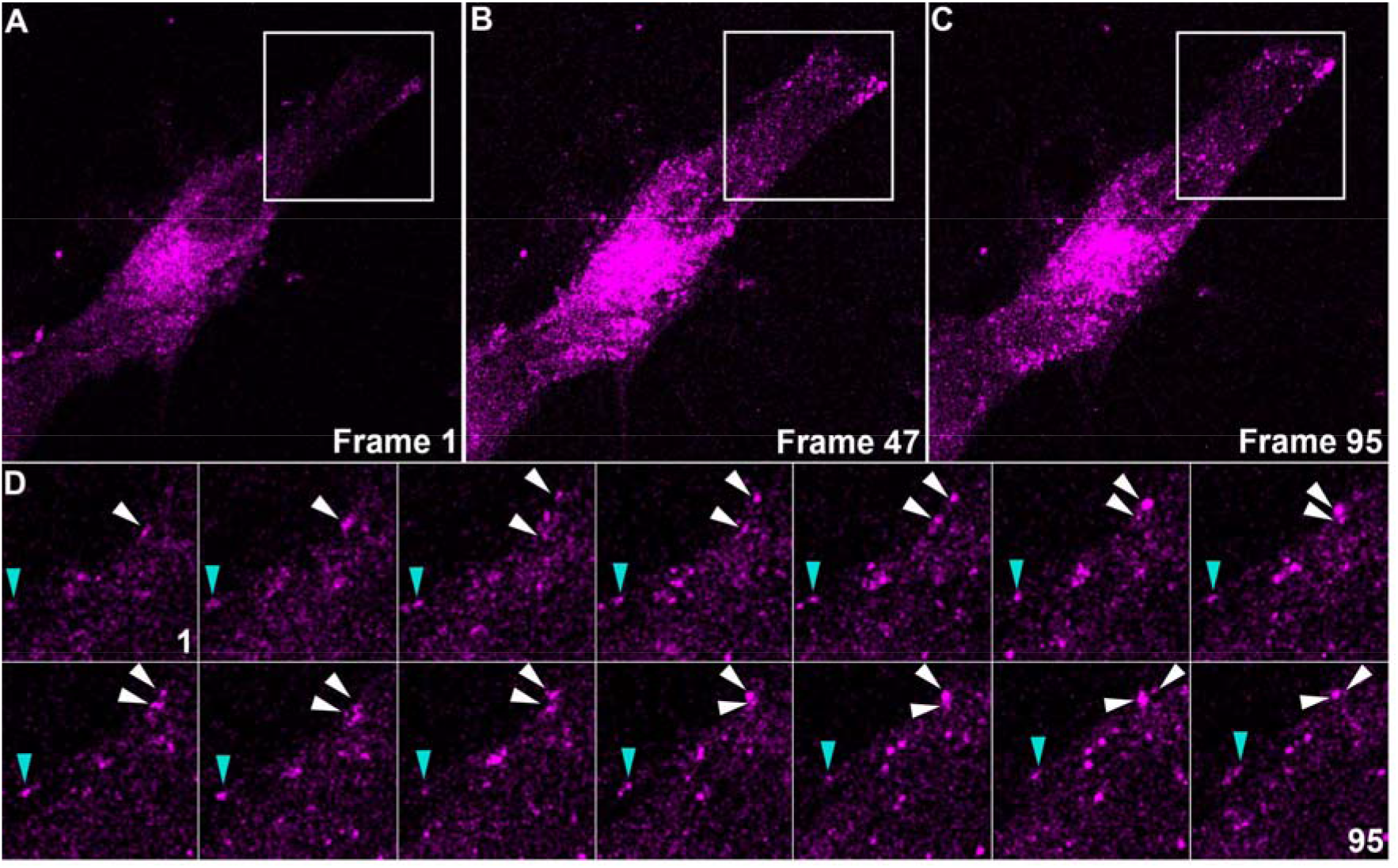
Optogenetic positioning of Rab11a endosomes to the cell periphery. A-C) Frames of live cell fluorescence imaging of optogenetic positioning of Rab11a endosomes. Light stimulation resulted in redistribution of Rab11a to the periphery of PtK_2_ cells transfected with Kif5a-GFP-CIBN, FuRed-CIBN-Rab11a and CRY2-Cluster-mCerulean. Cyan arrowhead = individual Rab11a endosome exhibiting slow trafficking to the cell periphery; white arrow heads = dynamic Rab11a endosomes trafficked to the cell periphery.

## Supplementary Movies

**Movie S1:** SBF-SEM of PtK_2_ grown on a R 2/2 carbon film gold support Quantifoil grid showing the fried egg morphology and abundant peripherally localised organelles.

**Movie S2:** Cryo-fLM confocal z-stack with Airyscan 2 detection EYFP-Mito-7 expressing PtK_2_ cell from ROI 1 (Fig 2A, C, C’) grown on +Carbon grids showing a thin cellular periphery.

**Movie S3:** Cryo-fLM confocal z-stack with Airyscan 2 detection EYFP-Mito-7 expressing PtK_2_ cell from ROI 2 (Fig 2A, D-H’’) grown on +Carbon grids demonstrating a thin cellular periphery.

**Movie S4:** Cryo-fLM confocal z-stack with Airyscan 2 detection EYFP-Mito-7 expressing PtK_2_ cell from ROI 3 (Fig S5A, C, C’) grown on +Carbon grids highlighting a z-stack of a cell too thick for cryo-ET analyses.

**Movie S5:** Higher-magnification (100X objective with 1.6X digital zoom) cryo-fLM confocal z-stack of EYFP-Mito-7 expressing PtK_2_ cell from ROI 3 (Fig S5A, C, C’) grown on +Carbon grids showing the thick cell periphery.

**Movie S6:** Cryo-fLM confocal z-stack with Airyscan 2 detection EYFP-Mito-7 expressing PtK_2_ cell from ROI 4 (Fig S5A, D, D’) grown on +Carbon grids highlighting a z-stack of a cell too thick for cryo-ET analyses.

**Movie S7:** Cryo-electron tomographic reconstruction of EGFP-DCX expressing PtK_2_ cell demonstrating abundant microtubules that align to the fluorescence signal from Fig S6B-H’’. Scale = 200 nm.

**Movie S8:** Live-cell fluorescence imaging of LAMP1-mCherry-CRY2 cell co-expressing KIF5a-GFP-CIBN and stimulated. Cell was imaged at 0.5s per frame.

**Movie S9:** Representative live-cell fluorescence time-lapse of a PtK_2_ cell transfected with Kif5a-GFP-CIBN, FuRed-CIBN-Rab11a and CRY2-Cluster-mCerulean. Light stimulation resulted in clustering of Rab11a endosomes and redistribution of Rab11a from predominantly perinuclear regions to the cell periphery. Cell was imaged at 0.5s per frame.

**Movie S10:** Cryo-electron tomographic reconstruction of stimulated LAMP1-mCherry-CRY2 and KIF5a-GFP-CIBN co-transfected PtK_2_ cell from ROI 1 demonstrating 3D organisation of the endolysosome from Fig 3G, G’. Scale = 200 nm.

**Movie S11:** Cryo-electron tomographic reconstruction of stimulated LAMP1-mCherry-CRY2 and KIF5a-GFP-CIBN co-transfected PtK_2_ cell from ROI 2 demonstrating 3D organisation of the endolysosome from Fig 3H, H’. Scale = 200 nm.

**Movie S12:** Cryo-electron tomographic reconstruction of stimulated LAMP1-mCherry-CRY2 and KIF5a-GFP-CIBN co-transfected PtK_2_ cell from ROI 3 demonstrating 3D organisation of the endolysosome from Fig 3I, I’. Scale = 200 nm.

## STAR Methods Table

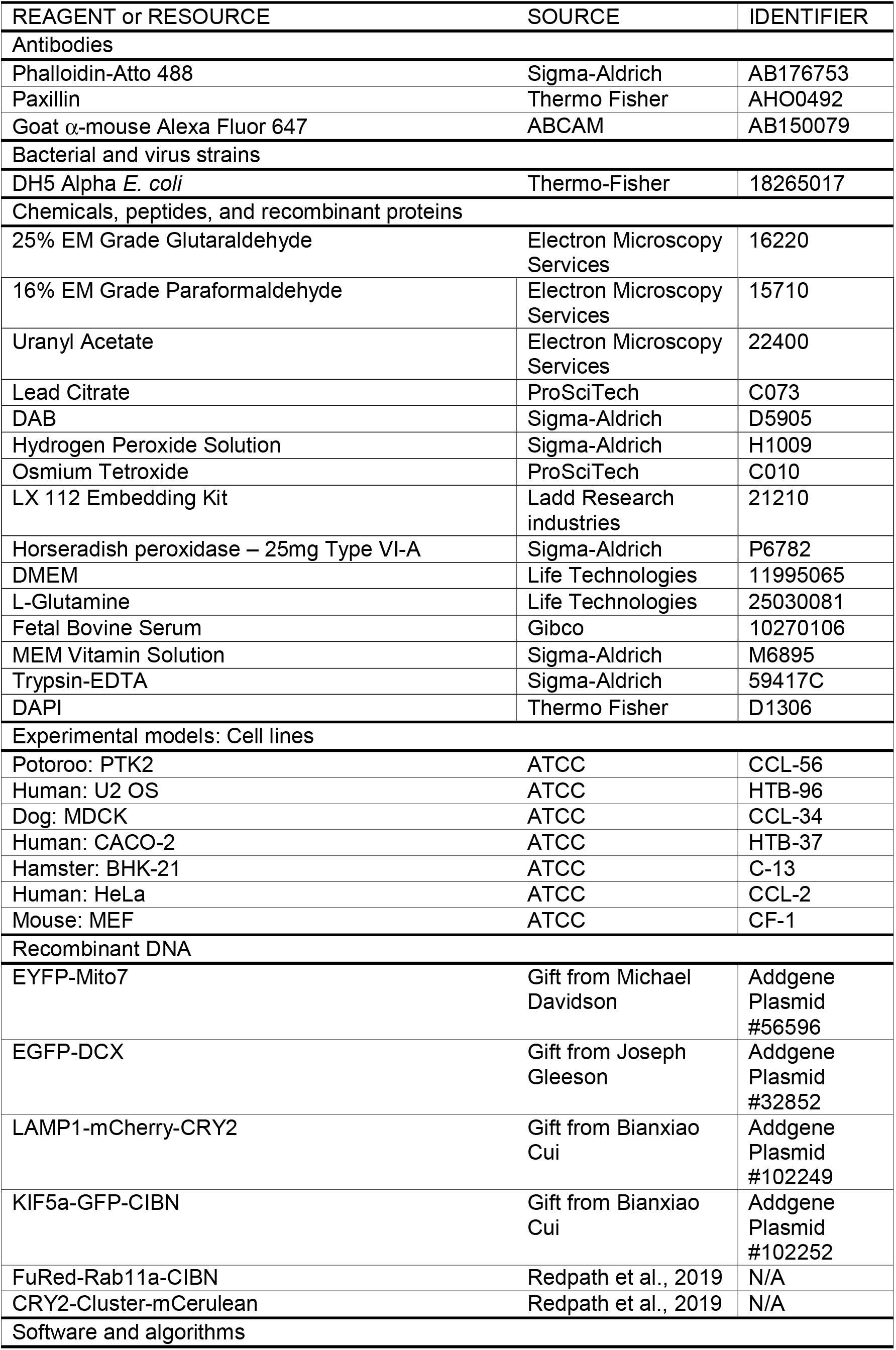
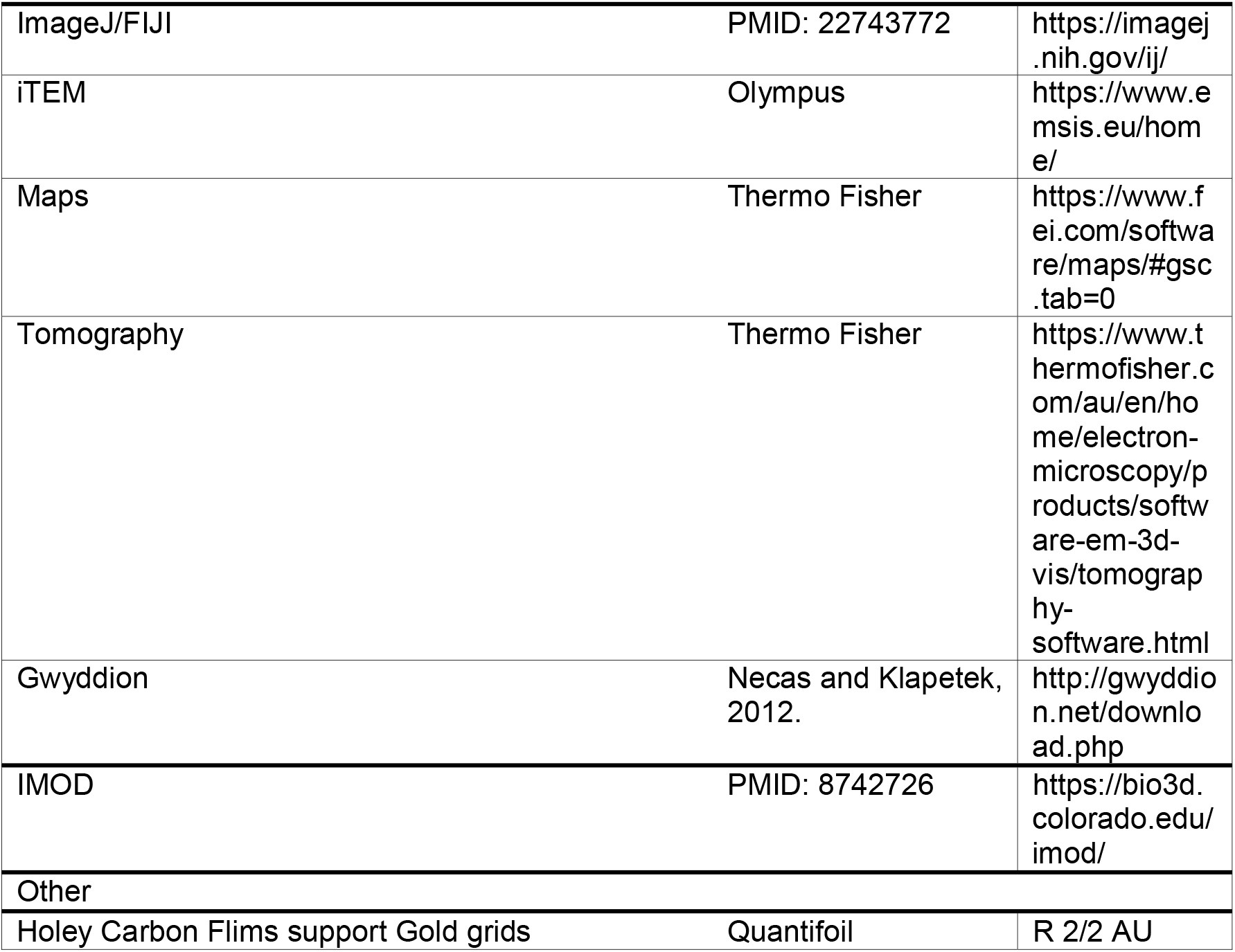

| REAGENT or RESOURCE | SOURCE | IDENTIFIER |
| --- | --- | --- |
| **Antibodies** |  |  |
| Phalloidin-Atto 488 | Sigma-Aldrich | AB176753 |
| Paxillin | Thermo Fisher | AHO0492 |
| Goat α-mouse Alexa Fluor 647 | ABCAM | AB150079 |
| **Bacterial and virus strains** |  |  |
| DH5 Alpha *E. coli* | Thermo-Fisher | 18265017 |
| **Chemicals, peptides, and recombinant proteins** |  |  |
| 25% EM Grade Glutaraldehyde | Electron Microscopy Services | 16220 |
| 16% EM Grade Paraformaldehyde | Electron Microscopy Services | 15710 |
| Uranyl Acetate | Electron Microscopy Services | 22400 |
| Lead Citrate | ProSciTech | C073 |
| DAB | Sigma-Aldrich | D5905 |
| Hydrogen Peroxide Solution | Sigma-Aldrich | H1009 |
| Osmium Tetroxide | ProSciTech | C010 |
| LX 112 Embedding Kit | Ladd Research industries | 21210 |
| Horseradish peroxidase – 25mg Type VI-A | Sigma-Aldrich | P6782 |
| DMEM | Life Technologies | 11995065 |
| L-Glutamine | Life Technologies | 25030081 |
| Fetal Bovine Serum | Gibco | 10270106 |
| MEM Vitamin Solution | Sigma-Aldrich | M6895 |
| Trypsin-EDTA | Sigma-Aldrich | 59417C |
| DAPI | Thermo Fisher | D1306 |
| **Experimental models: Cell lines** |  |  |
| Potoroo: PTK2 | ATCC | CCL-56 |
| Human: U2 OS | ATCC | HTB-96 |
| Dog: MDCK | ATCC | CCL-34 |
| Human: CACO-2 | ATCC | HTB-37 |
| Hamster: BHK-21 | ATCC | C-13 |
| Human: HeLa | ATCC | CCL-2 |
| Mouse: MEF | ATCC | CF-1 |
| **Recombinant DNA** |  |  |
| EYFP-Mito7 | Gift from Michael Davidson | Addgene Plasmid #56596 |
| EGFP-DCX | Gift from Joseph Gleeson | Addgene Plasmid #32852 |
| LAMP1-mCherry-CRY2 | Gift from Bianxiao Cui | Addgene Plasmid #102249 |
| KIF5a-GFP-CIBN | Gift from Bianxiao Cui | Addgene Plasmid #102252 |
| FuRed-Rab11a-CIBN | Redpath et al., 2019 | N/A |
| CRY2-Cluster-mCerulean | Redpath et al., 2019 | N/A |
| **Software and algorithms** |  |  |
| ImageJ/FIJI | PMID: 22743772 | https://imagej.nih.gov/ij/ |
| iTEM | Olympus | https://www.emsis.eu/home/ |
| Maps | Thermo Fisher | https://www.fei.com/software/maps/#gsc.tab=0 |
| Tomography | Thermo Fisher | https://www.thermofisher.com/au/en/home/electron-microscopy/products/software-em-3d-vis/tomography-software.html |
| Gwyddion | Necas and Klapetek, 2012. | http://gwyddion.net/download.php |
| IMOD | PMID: 8742726 | https://bio3d.colorado.edu/imod/ |
| Other |  |  |
| Holey Carbon Films support Gold grids | Quantifoil | R 2/2 AU |

